# A genome-wide genetic screen identified targets for destabilizing the parasitophorous vacuole of *Chlamydia trachomatis*

**DOI:** 10.1101/2024.08.02.606337

**Authors:** Mohammed Rizwan Babu Sait, Lana H. Jachmann, Milica Milivojevic, Celia Llorente-Sáez, Soniya Dhanjal, Fabian Schumacher, Sara Henriksson, Naga Venkata Gayathri Vegesna, Anastasiia Chaban, Partha Mohanty, Magnus Ölander, Samada Muraleedharan, Sepideh Farmand Azadeh, Burkhard Kleuser, Bernhard Schmierer, Barbara S. Sixt

## Abstract

The bacterial pathogen *Chlamydia trachomatis* employs the effector CpoS to suppress a host defense response that aborts intracellular bacterial growth by inducing host cell death. While conducting a CRISPR knock-out screen for genes contributing to this response, we uncovered a mutant deficient for CpoS to display a markedly increased reliance on host cellular ceramide synthesis, compensating for its diminished ability to acquire sphingolipids via modulating membrane trafficking. Employing the power of the just recently established molecular genetic toolbox for *Chlamydia*, we developed an innovative microscopic reporter system that revealed the mutant to thrive in unstable parasitophorous vacuoles (inclusions), characterized initially by the release of individual bacteria from otherwise intact-appearing vacuoles. CpoS-deficient inclusions were further destabilized by disruptions in ceramide synthesis, while supplementation of sphingoid bases stabilized them, also preventing the defensive host cell death response. Notably, early inclusion destabilization, achieved by simultaneous disruption of two transport routes, caused infection clearance without damaging the host cells. Overall, this study highlights the inclusion’s role as a refuge, demonstrates CpoS to maintain inclusion integrity by ensuring sphingolipid supply, and provides directions for a future therapeutic exploitation.

**SIGNIFICANCE:** A wide range of clinically significant microbes evolved to hide from the intrinsic defenses of their host cells by thriving within membrane-enclosed pathogen-containing vacuoles. This raises the intriguing possibility that such vacuoles could be targeted therapeutically. The bacterial pathogen *Chlamydia trachomatis* could be an exceptionally well-suited target for such innovative medicines given its medical importance and strict dependence on host cells. However, progress has been stalled by the lack of sensitive tools for detecting inclusion damage. Here, we resolved this major technical roadblock and uncovered the pathogen to employ the secreted effector CpoS, a modulator of membrane trafficking, to stabilize its vacuole by ensuring adequate sphingolipid supply. These methodological advances and mechanistic insights should promote the development of vacuole-destabilizing therapeutics.

## INTRODUCTION

To flourish within human cells, intracellular pathogens must adeptly circumvent their host cells’ defenses. Targeting this intricate interplay between host cell-autonomous immunity and pathogen evasion strategies could provide novel therapeutic avenues. Yet, our insufficient understanding of the underlying mechanisms remains a major roadblock. Here, we report on how the bacterial pathogen *Chlamydia trachomatis* preserves its parasitophorous vacuole to hide from its host cell, and we uncover possible targets for a deliberate vacuole destabilization.

Initially identified as the cause of trachoma, a blinding ocular disease that remains a significant public health concern in more than 40 countries ^1^, *C. trachomatis* is now recognized as the major bacterial agent of sexually transmitted infections, which can inflict enduring harm to reproductive tissues and accounts for over 130 million cases annually ^2,3^. The pathogen exhibits a high dependency on epithelial host cells due to its obligate intracellular lifestyle and developmental biology, characterized by an alternation between two distinct forms: the replicative reticulate body (RB) and the infectious elementary body (EB) ^4^. Upon infiltrating a host cell, the EB settles within a membrane-bound vacuole, the inclusion, where it differentiates into the RB form, which then replicates and enlarges the vacuole. Around 24 hours post-infection (hpi), the RBs begin reverting back into EBs, which are released around 48-72 hpi via host cell lysis or inclusion extrusion ^5^.

*C. trachomatis* employs an arsenal of secreted effector proteins to modulate host cell functions and shape its intracellular niche ^6^. This includes a unique class of effectors, the inclusion membrane proteins (Incs), which the pathogen inserts into the membrane of its vacuole ^7^. The recent introduction of tools enabling the molecular genetic manipulation of the pathogen revolutionized our capacity to decipher the critical functions of these effectors ^8^. Significantly, these novel approaches revealed the Inc CpoS to suppress host cellular immune surveillance ^9,10^.

Infections with CpoS-deficient mutants lead to a potent activation of the STING/TBK1/IRF3 immune signaling pathway, resulting in increased type I interferon (IFN) production and induction of IFN-stimulated genes ^9^. The host cells then also succumb to death prematurely, effectively disrupting bacterial replication and EB formation and causing an accelerated clearance from the genital tract of experimentally infected mice ^9,10^. While the defensive cell death response requires host cellular protein synthesis ^9^, suggesting it is actively triggered by the host cells, it is not dependent on a functional type I IFN response ^9^, thus cannot be considered a direct result of the enhanced IFN signaling. Furthermore, while CpoS’ interactions with host Rab GTPases are crucial in the effector’s ability to modulate membrane trafficking and dampen type I IFN responses, reinstating Rab interactions proofed insufficient in preventing premature host cell death ^11^.

The nature of the host defense program inhibited by CpoS thus remained unclear. Setting out to unravel its mechanistic basis through an unbiased genome-wide genetic screen, we made unexpected discoveries, which highlighted a possible contribution of sphingolipid imbalances to the cell death, as well as the crucial role of these lipids in preserving the integrity of *Chlamydia*’s parasitophorous vacuole, which shields the pathogen from host cellular defenses. Overall, these observations should facilitate the future therapeutic targeting of the *Chlamydia* inclusion.

## RESULTS

### A CRISPR screen identified deficiencies protective against *Chlamydia*-induced cytotoxicity

To identify host genes crucial for the execution of the defensive cell death response inhibited by CpoS, we conducted a CRISPR/Cas9 knockout screen for host gene deficiencies that protect cells from death induced by a *C. trachomatis cpoS* null mutant (Fig 1A and S1A). Moreover, to aid in pinpointing those genes that contribute to this response specifically, but not to *Chlamydia*-induced cytotoxicity or *Chlamydia* infection generally, we concurrently screened for host gene deficiencies protecting cells from the wild-type strain of *C. trachomatis*. To this end, we transduced HeLa cells, a human cervical epithelial cell line, with a genome-wide knockout library. We then infected the cells with *C. trachomatis* L2/434/Bu, either wildtype (CTL2) or carrying an insertional disruption of *cpoS* (CTL2-*cpoS*::*cat*). Later, we collected cells resistant to infection-mediated killing, that is, cells resistant to late-stage lytic death in the case of CTL2 or premature death in the case of CTL2-*cpoS*::*cat*. Hence, we included four conditions: uninfected cells and cells infected with CTL2-*cpoS*::*cat* at 30 hpi (UI30h and KO30h), and uninfected cells and cells infected with CTL2 at 60 hpi (UI60h and WT60h). The screen was performed in two independent replicates (R1+R2).

**Figure 1.**
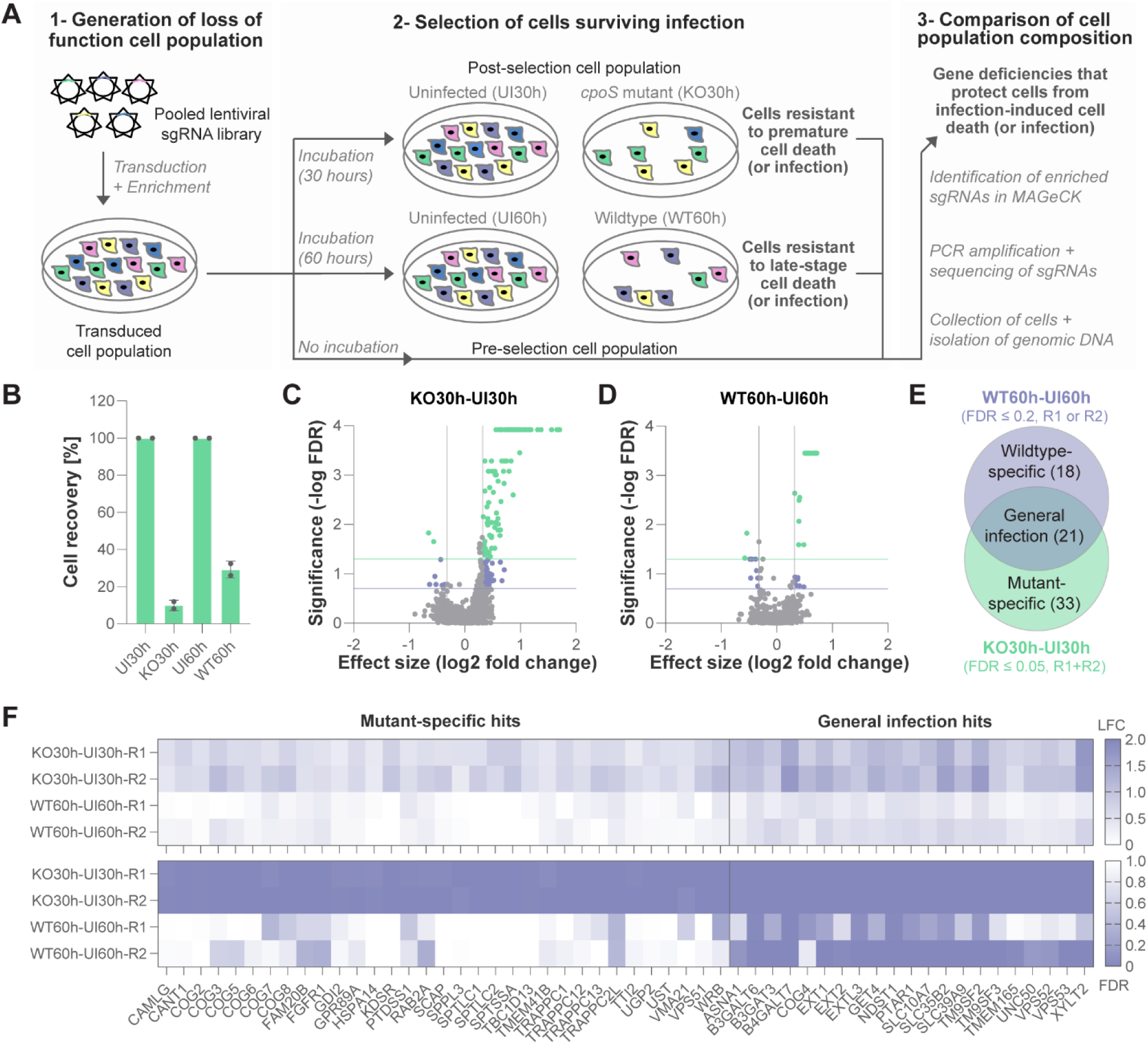
A CRISPR screen identified deficiencies protective against *Chlamydia*-induced cytotoxicity. **(A)** CRISPR screening procedure. See Fig S1A, for a more detailed illustration of the selection. **(B)** Proportion of cells (relative to the respective uninfected controls) that could be recovered at the sampling times (n=2 (R1, R2), mean±SD). **(C-D)** Volcano plots displaying genes with depleted or enriched sgRNAs in cultures infected with CTL2-*cpoS*::*cat* (KO30h vs UI30h, C) or CTL2 (WT60h vs UI60h, D). Shown are data for one replicate. Marked in green, FDR ≤ 0.05 (-log FDR ≥ 1.3) and FC ≤ 0.8 or ≥ 1.25 (log2 FC ≤ -0.32 or ≥ 0.32); marked in lilac, FDR ≤ 0.2 (-log FDR ≥ 0.7) and FC ≤ 0.8 or ≥ 1.25 (log2 FC ≤ -0.32 or ≥ 0.32). **(E)** Venn diagram illustrating the selection of screening hits. **(F)** Heatmap displaying enrichment FDR and log2 fold change (LFC) of the screening hits for the indicated comparisons and replicates.

Our initial aim was to collect the cells when about 90% had died, ensuring a robust enrichment of cells with reduced susceptibility to infection-mediated killing. In the screen, this was indeed achieved for cells infected with CTL2-*cpoS*::*cat* when collected at 30 hpi (Fig 1B and S1A). However, cells infected with CTL2 had to be collected at 60 hpi, when only about 75% had perished, because delaying the collection further would have led to significant loss during harvesting, given the fragility of cells containing large inclusions.

Upon concluding the selection experiments, we sequenced the sgRNAs. This confirmed a good library representation, with over 99.9% of all library sgRNAs detected in all samples (Fig S1B), and no drastic shifts in population composition (Fig S1C). We then identified sgRNAs that were positively selected, *i.e.*, sgRNAs that showed significant enrichment in infected compared to uninfected samples in pairwise comparisons (Table S1). Those sgRNAs could be assumed to target genes that normally promote infection-mediated cell killing. By setting a false discovery rate (FDR) of ≤ 0.05 and an average fold change (FC) of ≥ 1.25 as cutoffs, we found 54 genes with enriched sgRNAs in KO30h vs UI30h in both R1 and R2 (Fig 1C and S1D). Conversely, sgRNAs for only three genes showed enrichment in WT60h vs UI60h in both R1 and R2 (Fig 1D and S1E), aligning with the weaker selection strength achieved for this strain. To get a more comprehensive overview of genes contributing to *Chlamydia*-induced cytotoxicity or infection in general, we also considered sgRNAs enriched in WT60h vs UI60h at FDR ≤ 0.2. Indeed, we observed these to partially overlap with those found enriched in KO30h vs UI30h, and we concluded the overlapping sgRNAs to more likely target genes with critical roles during infection rather than in the *cpoS* mutant triggered cell death response.

Consequently, we selected hits as follows (Fig 1E-F, Table S2): We classified genes targeted by sgRNAs enriched in both KO30h vs UI30h (FDR ≤ 0.05 and FC ≥ 1.25, R1+R2) and WT60h vs UI60h (FDR ≤ 0.2 and FC ≥ 1.25, at least one replicate) as “general infection hits”, resulting in 21 genes. We identified “mutant-specific hits” as genes whose sgRNAs were enriched in KO30h vs UI30h (FDR ≤ 0.05 and FC ≥ 1.25, R1+R2) but not WT60h vs UI60h (FDR ≤ 0.2, any replicate), yielding 33 genes. Although genes with enriched sgRNAs in WT60h vs UI60h (FDR ≤ 0.2 and FC ≥ 1.25, any replicate), but not KO30h vs UI30h (FDR ≤ 0.05, R1+R2), could be considered as “wildtype-specific hits”, these would be of low confidence and were not further evaluated here.

### Deficiencies in heparan sulfate proteoglycan synthesis provided general protection against *C. trachomatis* L2

To comprehend the biological roles of the genes targeted by positively selected sgRNAs, we conducted a functional enrichment analysis in g:Profiler ^12^ using the Reactome ^13^ pathway database. This revealed that the “general infection hits” were enriched for genes involved in proteoglycan biosynthesis and intra-Golgi trafficking (Fig 2A-B), while the “mutant-specific hits” were enriched for genes involved in various vesicular transport processes and sphingolipid metabolism (Fig 2C-D). However, only a small subset of the genes in these gene lists was represented in the functional terms highlighted by the g:Profiler analysis.

**Figure 2.**
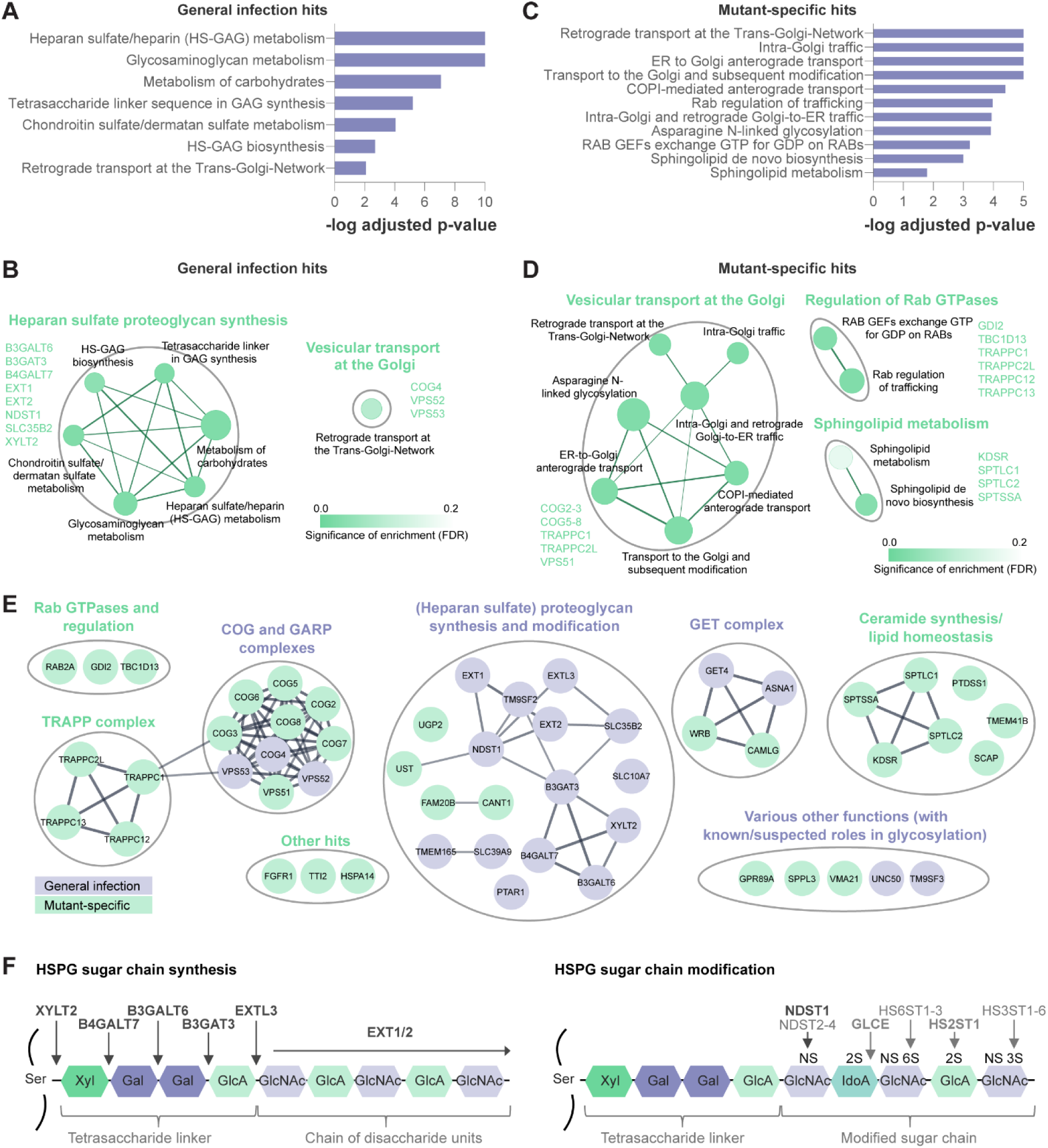
Deficiencies in heparan sulfate proteoglycan synthesis provided general protection against *C. trachomatis* L2. (A-D) Functional enrichment analysis of “general infection hits” (A-B) and “mutant-specific hits” (C-D) conducted in g:Profiler ^12^ using the Reactome pathway database ^13^. Bar plots display enrichment p-values for all significantly (p ≤ 0.05) enriched pathway terms (A, C). Cytoscape enrichment maps ^41,42^ display the relationship between significantly enriched pathway terms (B, D). Nodes represent pathways (node size indicates the number of genes in each pathway, node color the significance of the enrichment), and edges their similarity (edge widths indicate the size of similarity). Connected pathways were grouped. For each group, all genes found enriched are indicated. **(E)** STRING interaction network displaying interactions among and between “general infection hits” (lilac nodes) and “mutant-specific hits” (green nodes), refined by manual grouping and annotations. Mixed groups were deemed to represent biological processes of general importance for infection (lilac group labels), while groups containing only “mutant-specific hits” were considered to have specific importance during infection with the *cpoS* mutant (green group labels). **(F)** Deficiencies in the HSPG biosynthetic pathway conferred general protection against *C. trachomatis* L2. HSPG synthesis starts with the attachment of a tetrasaccharide linker composed of xylose (Xyl), galactose (Gal), and glucuronic acid (GlcA) to specific serine residues on the core protein (catalyzed by XYLT2, B4GALT7, B4GALT6, B3GAT3), followed by elongation of the sugar chain (catalyzed by EXTL3, EXT1, EXT2) through addition of disaccharide units composed of N-acetylglucosamine (GlcNAc) and glucuronic acid (GlcA). Subsequently, NDST mediates the sulfation (NS, N-sulfation) and de-acetylation of specific GlcNAc residues, GLCE induces the epimerization of GlcA to iduronic acid (IdoA), and the enzymes HS2ST, HS6ST, and HS3ST mediate the attachment of sulfate groups to specific saccharide residues (2S, 2-O-sulfation; 6S, 6-O-sulfation; 3S, 3-O-sulfation). Enzymes encoded by genes whose deficiency protected cells from infected-induced toxicity according to the CRISPR screen are labeled in bold dark gray (or bold light gray if found with lower confidence).

To gain a deeper understanding of the major biological processes represented by our hits, we utilized the STRING database ^14^ to display a protein interaction network, which we then further refined by manual grouping and annotation based on literature (Fig 2E, Table S3). This in-depth analysis highlighted very concrete molecular pathways and complexes, as well as significant functional overlaps between the “general infection hits” and “mutant-specific hits”. Indeed, the two gene lists contained constituents of the same molecular machineries and processes, such as heparan sulfate proteoglycan (HSPG) biosynthesis, conserved oligomeric Golgi (COG) complex, Golgi-associated retrograde protein (GARP) complex, and guided entry of tail-anchored proteins (GET) pathway. Again, given the overlap, we concluded that these machineries and pathways were more likely to have fundamental roles during *C. trachomatis* L2 infection rather than specific roles in the defense response inhibited by CpoS.

Interestingly, genes with key or auxiliary functions in the synthesis of HSPGs, cell surface molecules known to facilitate host cell attachment and invasion by *C. trachomatis* L2 ^15–17^, were especially dominant among our hits (Fig 2E-F, Table S2-S3). Specifically, this included genes encoding the four enzymes required for the synthesis of the tetrasaccharide linker common to both HSPGs and chondroitin sulfate proteoglycans (XYLT2, B4GALT7, B3GALT6, B3GAT3) ^18^, the enzymes involved in sugar chain elongation specific for HSPGs (EXTL3, EXT1, EXT2) ^18^, proteins involved in further glycan chain modifications (NDST1, FAM20B, UST; with lower confidence also GLCE and HS2ST1) ^18,19^, proteins that mediate the synthesis or transport of necessary precursors or the removal of side products (SLC35B2, UGP2, CANT1) ^18,20,21^, and other proteins shown to be important for proteoglycan synthesis due to their role in maintaining a functional glycosylation or glycan-modifying machinery (*e.g.*, TM9SF2, TMEM165, SLC10A7, SLC39A9, PTAR1) ^22–29^. Moreover, several of our other hits may have roles at least in part also associated with HSPG synthesis. For example, the GET pathway, which mediates membrane insertion of tail-anchored proteins ^30^, and the complexes COG and GARP, which have roles in membrane trafficking ^31,32^, may regulate the localization of proteins involved in HSPG synthesis or modification. Indeed, deficiencies in COG and GARP have been shown to cause mislocalization and destabilization of proteins of the glycosylation machinery at the Golgi ^33,34^. Moreover, deficiencies in SPPL3, GRP89A, and VMA21 have also been linked to defects in Golgi homeostasis and/or glycosylation ^35–39^, and sgRNAs targeting UNC50 and TM9SF3 have in CRISPR screens frequently been found co-enriched with those targeting HSPG biosynthesis (*^e.g.,^* ^26,29,40^).

In conclusion, this first mining of the data revealed that our screening approach had pointed out very concrete molecular processes and machineries to have important roles during infection with *C. trachomatis* L2. Critically, it underscored the significance of HSPGs, which validated the performance of the screen and enabled us to focus our attention on other processes offering specific insights into the function of CpoS and the defense response it inhibits.

### Deficiencies in ceramide synthesis provided specific protection against the *cpoS* mutant

Among the host gene deficiencies providing specific protection against the *cpoS* mutant (Fig 2E, Tables S2-S3), we were surprised to find none in genes known to act in pathogen recognition, immune signaling, or regulated cell death. Instead, these deficiencies impacted components of the multi-subunit tethering complex TRAPP (TRAPPC1, TRAPPC2L, TRAPPC12, TRAPPC13) and the Rab GTPase RAB2A, both involved in trafficking between ER and Golgi ^43,44^, as well as regulators of Rab protein function (GDI2, TBC1D13) ^44,45^. Intriguingly, deficiencies in the *de novo* synthesis of ceramides also conferred specific protection. This included deficiencies in any of the three subunits of serine palmitoyltransferase (SPT, namely SPTLC1, SPTLC2, SPTSSA), which catalyzes the first enzymatic step ^46^, and deficiency in 3-ketodihydrosphingosine reductase (KDSR), responsible for the second enzymatic step ^46^ (Fig 3A).

**Figure 3.**
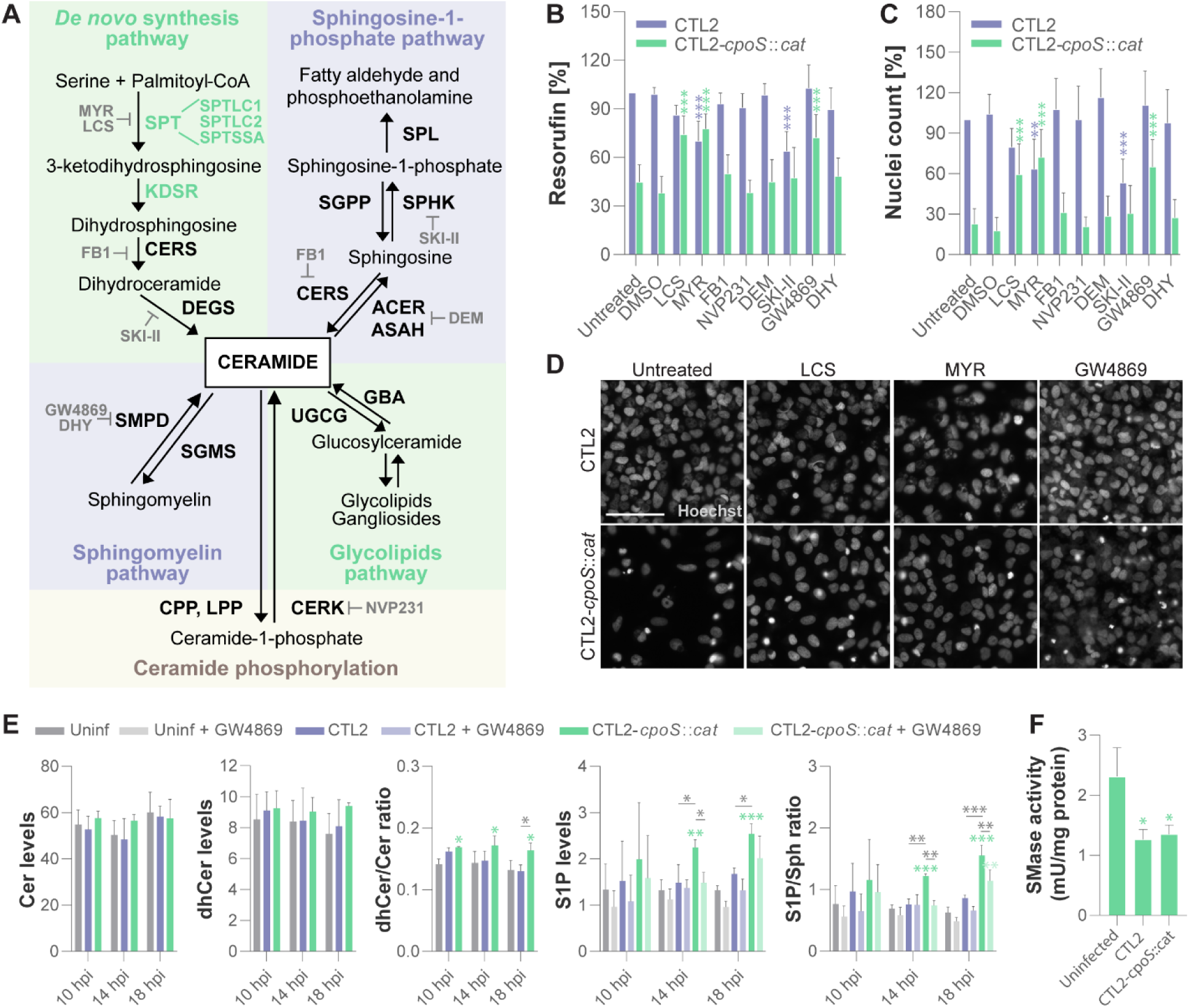
Deficiencies in ceramide synthesis provided specific protection against the *cpoS* mutant. **(A)** Illustration of pathways in ceramide synthesis and metabolism. Enzymes whose deficiency protected cells from *cpoS* mutant-induced toxicity in the screen are highlighted in bold green, pharmacologic inhibitors in bold gray. ACER/ASAH, ceramidase; CERK, ceramide kinase; CERS, ceramide synthase; CPP, ceramide-1-phosphate phosphatase; DEGS, dihydroceramide desaturase; DEM, D-erythro-MAPP; DHY, desipramine hydrochloride; FB1, fumonisin B1; GBA, glucosylceramidase; KDSR, 3-ketodihydrosphingosine reductase; LCS, L-cycloserine; LPP, lipid phosphate phosphatase; MYR, myriocin; SGMS, sphingomyelin synthase; SGPP, sphingosine-1-phosphate phosphatase; SMPD, sphingomyelinase; SPHK, sphingosine kinase; SPL, sphingosine phosphate lyase; SPT, serine palmitoyl transferase; UGCG, UDP-glucose ceramide glycosyltransferase. **(B-D)** MYR, LCS, and GW4869 protected cells from *cpoS* mutant-induced death. HeLa cells were treated with the indicated inhibitors (LCS, 125 µM; MYR, 1 µM; FB1, 25 µM; NVP231, 0.625 µM; DEM, 25 µM; SKI-II, 6.25 µM; GW4869, 12.5 µM; DHY, 12.5 µM) or solvent only (DMSO), and were parallelly infected with the indicated strains (4 IFU/cell). Resorufin fluorescence (B) and nuclei count (C) at 25.5 hpi are displayed normalized to a CTL2-infected untreated control (mean±SD, n=7, one-way ANOVA with Dunnett’s; for each strain, indicated are significant differences compared to the untreated control). (D) Representative images (scale = 100 µm). **(E)** Quantification of selected sphingolipid metabolites. HeLa cells were infected with the indicated strains in the presence or absence of GW4869 (12.5 µM). At the indicated time points, cell extracts were prepared, and the indicated lipids were quantified by LC-MS/MS. Shown are selected metabolite levels (expressed as “fmol/pmol total sphingolipids”) or ratios (mean±SD, n=3, one-way ANOVA with Tukey’s; if not specified otherwise, for each time point, indicated are significant differences compared to uninfected untreated cells; Cer, ceramides; dhCer, dihydroceramides; S1P, sphingosine-1-phosphate; Sph, sphingosine). Full data are presented in Fig S3. **(F)** Neutral sphingomyelinase activity detected in cell lysates prepared at 18 hpi from HeLa cells infected with the indicated strains (5 IFU/cell; mean±SD, n=3, one-way ANOVA with Tukey’s; indicated are significant differences compared to uninfected cells).

Ceramides and derived metabolites are signaling molecules with key roles in cellular life/death decisions ^47,48^. Therefore, it seemed plausible that a deficiency in *cpoS* could instigate host cell death by disturbing metabolite balance, for instance resulting in an accumulation of ceramides or an increase in the dihydroceramide/ceramide ratio, effects that could potentially be mitigated by impairments in the ceramide synthesis pathway. To explore this hypothesis, we assessed whether inhibiting enzymes involved in ceramide metabolism could protect cells against *cpoS* mutant-induced death. We specifically tested inhibitors of SPT [myriocin (MYR) and L-cycloserine (LCS)], ceramide synthase [fumonisin B1 (FB1)], neutral sphingomyelinase [GW4869], acid sphingomyelinase [desipramine hydrochloride (DHY)], alkaline ceramidase [D-erythro-MAPP (DEM)], sphingosine kinase [SKI-II], and ceramide kinase [NVP231] (Fig 3A). HeLa cells were treated with the respective inhibitors at for uninfected cells non-toxic concentrations (Fig S2) and simultaneously infected with either CTL2 or CTL2-*cpoS*::*cat*. Host cell viability was quantified at 24 hpi based on the ability of viable cells to convert resazurin into fluorescent resorufin (Fig 3B) and the number of remaining cell nuclei (Fig 3C-D). The SPT inhibitors effectively protected cells from death induced by the *cpoS* mutant. Additionally, we observed reduced cell death also in cells treated with GW4869. Interestingly, SKI-II, which besides sphingosine kinase also inhibits dihydroceramide desaturase and may thereby increases the dihydroceramide/ceramide ratio ^49^, had the opposite effect, inducing cell death in cells infected with CTL2 (Fig 3B-C).

In summary, these findings indicated that deficiencies in or the inhibition of ceramide-producing enzymatic reactions can protect cells from *cpoS* mutant-induced death. Moreover, they revealed that certain modulations in the sphingolipid metabolism can indeed induce the death of *Chlamydia*-infected cells.

### CpoS deficiency caused alterations in the levels of certain sphingolipid metabolites

To determine if the premature host cell death during infection with the *cpoS* mutant could be due to an intracellular build-up of ceramides or dihydroceramides or other alterations in sphingolipid levels, we employed mass spectrometry to quantify the intracellular levels of 31 sphingolipid metabolites, encompassing a variety of common sphingoid bases, dihydroceramides, ceramides, dihydrosphingomyelins, sphingomyelins, hexosylceramides, and lactosylceramides. Our focus was on the early stages of infection, both before and at the onset of premature host cell death. Overall, we did not detect any remarkable differences in the metabolite levels between uninfected cells and those infected with CTL2, and most metabolite levels remained similar also during infection with CTL2-*cpoS*::*cat* (Fig S3A-B). Interestingly, we did not observe a rise in bulk ceramide or dihydroceramide levels (Fig 3E). However, we did indeed detect a significant increase in the dihydroceramide/ceramide ratio during infection with CTL2-*cpoS*::*cat*, as well as notable increases in the levels of sphingosine-1-phosphate (S1P) and the S1P/sphingosine ratio (Fig 3E). The administration of the neutral sphingomyelinase inhibitor GW4869 partially mitigated the latter effects, suggesting that the elevated levels of S1P might at least in part be derived from the hydrolysis of sphingomyelins into ceramides and their further conversion to sphingosine and S1P. Unexpectedly, we found neutral sphingomyelinase activity to be reduced during infection, and to be similar in cells infected with either CTL2 or CTL2-*cpoS*::*cat* (Fig 3F).

In summary, our findings indicate significant changes in the occurrence of certain sphingolipid metabolites in cells infected with the *cpoS* mutant. This includes an increase in the dihydroceramide/ceramide ratio, an alteration known to induce cell death ^48^. Moreover, while neither bulk levels of ceramides nor neutral sphingomyelinase activity were increased, the elevated levels of S1P may imply an activation of sphingomyelinase and the production of pro-death ceramides to occur in a more localized manner at specific subcellular locations.

### The *cpoS* mutant displayed an increased dependence on *de novo* ceramide synthesis

Given that MYR, LCS, and GW4869 partially blocked *cpoS* mutant-induced premature cell death, we hypothesized they could also restore the intracellular growth of the mutant. Hence, we treated HeLa cells and concurrently infected them with CTL2 or CTL2-*cpoS*::*cat*. At 24 hpi, we detected bacterial inclusions microscopically (Fig 4A-B). As expected, due to the host cell death, we saw significantly fewer inclusions for CTL2-*cpoS*::*cat* compared to CTL2 in the absence of inhibitors. Moreover, all three inhibitors caused minor reductions in the number and size of CTL2 inclusions. Unexpectedly, LCS and MYR not only failed to restore the growth of CTL2-*cpoS*::*cat* but almost entirely abrogated it, whereas in the presence of GW4869, the number of inclusions remained similar to that in the untreated control. A comparison of growth inhibition across a range of inhibitor concentrations confirmed that the *cpoS* mutant was considerably more reliant on SPT function, while both strains appeared to respond similarly to GW4869 (Fig 4C).

**Figure 4.**
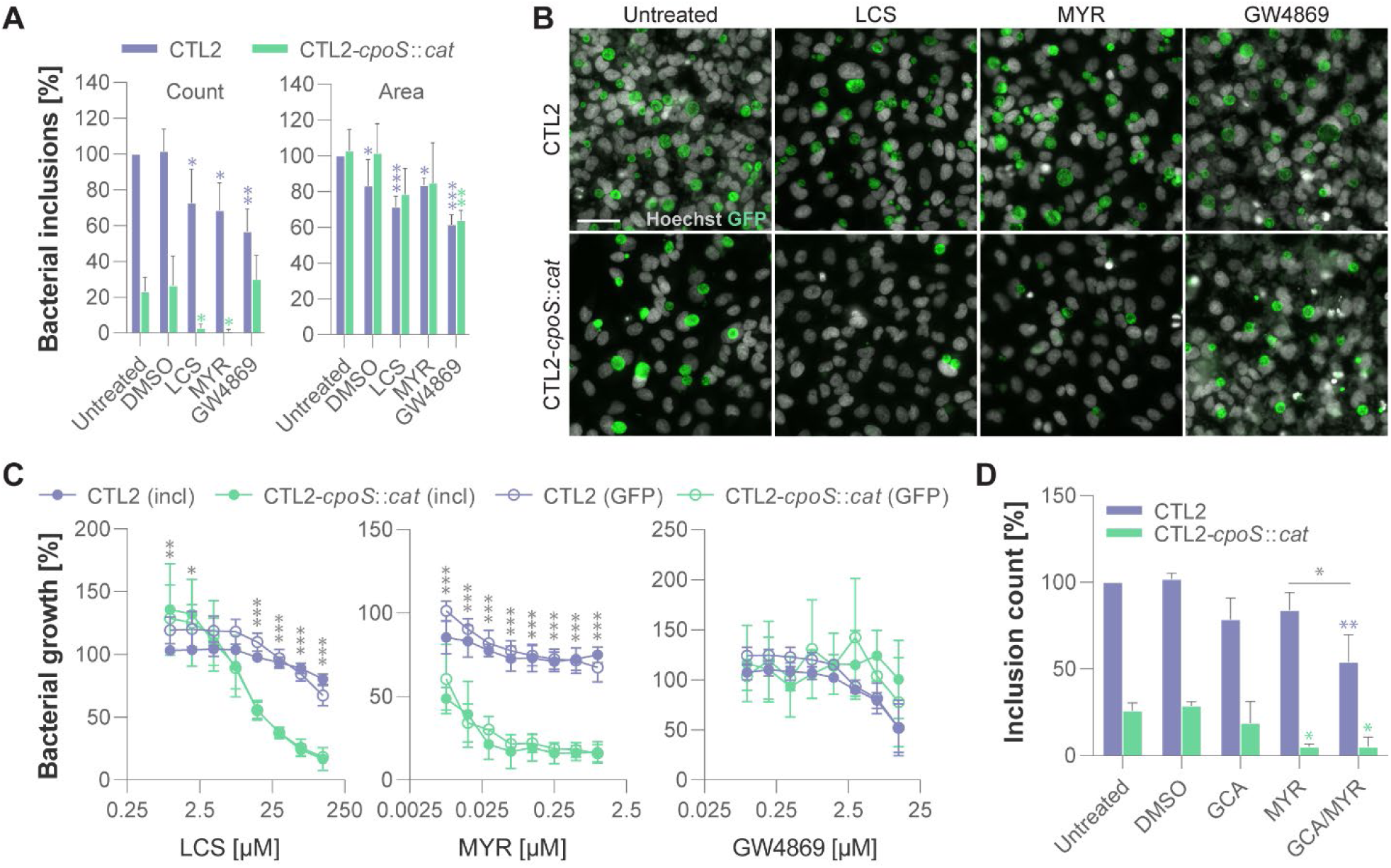
The *cpoS* mutant displayed an increased dependence on *de novo* ceramide synthesis. (A-B) MYR and LCS, but not GW4869, abrogated the growth of the *cpoS* mutant. HeLa cells were treated with the indicated inhibitors (LCS, 125 µM; MYR, 1 µM; GW4869, 12.5 µM) or DMSO only, and parallelly infected with GFP-expressing derivatives of the indicated strains (1 IFU/cell). Bacterial inclusions were detected at 26.5 hpi. (A) Data displayed as percentage relative to inclusion count and average inclusion area for CTL2 in the absence of inhibitors (mean±SD, n=4, one-way ANOVA with Dunnett’s; for each strain, indicated are significant differences compared to the untreated control). (B) Representative images (scale = 50 µm). **(C)** Differential susceptibility of *cpoS* mutant and wild-type bacteria to SPT inhibition. HeLa cells were treated with the indicated inhibitors and parallelly infected with GFP-expressing derivatives of the indicated strains (1 IFU/cell). GFP fluorescence (GFP) and the number of inclusions (incl) at 25.5 hpi are displayed relative to values detected for the strain in the absence of inhibitors (mean±SD, n=3-4, 2-way ANOVA with Sidak’s; for each concentration, indicated are significant differences in inclusion counts between the two strains). **(D)** Blockade in membrane trafficking sensitized CTL2 to the growth-inhibitory action of MYR. HeLa cells were treated with the indicated inhibitors (MYR, 0.5 µM; GCA, 5 µM) and parallelly infected with GFP-expressing derivatives of the indicated strains (2 IFU/cell). Inclusion numbers at 20 hpi are displayed relative to CTL2 in the absence of inhibitors (mean±SD, n=2, one-way ANOVA with Tukey’s; if not specified otherwise, for each strain, indicated are significant differences compared to the untreated control).

Ceramides serve a dual role. They are signaling lipids but also precursors for the synthesis of complex sphingolipids, such as sphingomyelin, which *C. trachomatis* requires for its growth ^50^. The bacteria can acquire sphingolipids in two ways, by recruiting the host ceramide transfer protein CERT to acquire ceramides then converted to complex sphingolipids at the inclusion ^51^, and by hijacking sphingolipid-laden membrane vesicles and directing them to the inclusion ^52^. We previously reported CERT recruitment to be unaffected by CpoS-deficiency, while the vesicular transport of sphingomyelin to inclusions was compromised ^11^. Hence, the mutant’s heightened dependence on *de novo* ceramide synthesis could be attributed to its reduced capacity to utilize the available sphingolipid pool. Indeed, when we inhibited vesicular transport of Golgi-derived sphingomyelin to the inclusion, using golgicide A ^51^, the dependence of CTL2 on host *de novo* ceramide synthesis intensified (Fig 4D). Interestingly, the CRISPR screening data also suggested a trend where CERT-deficiency enhanced the susceptibility of cells to CTL2-induced cytotoxicity and protected cells from *cpoS* mutant-induced cell death (Fig S4A). An attempt to recapitulate this finding by exposing infected cells to the CERT-inhibitory ceramide analog HPA-12 revealed both strains to display similar sensitivity to its growth-inhibitory action (Fig S4B), which, however, could be due to HPA-12 blocking *Chlamydia* growth also in CERT-independent manners ^53^.

In summary, our research indicated that CpoS deficiency heightens the bacteria’s reliance on host cellular ceramide synthesis. Consequently, the protection against cell death, which is observed when SPT is inhibited, can potentially be interpreted as an indirect outcome of inhibiting bacterial growth. Conversely, GW4869 appears to block cell death via a different mechanism.

### A novel reporter for inclusion damage revealed instability of CpoS-deficient inclusions

Having uncovered that SPT inhibition blocks *cpoS* mutant-induced cell death indirectly, we sought to obtain a deeper mechanistic understanding of the ways by which sphingolipid deprivation can curb the growth of the mutant. It’s worth noting that CpoS-deficiency has been proposed to cause premature inclusion rupture ^10^. Moreover, prior work has suggested that both SPT inhibition and disruptions in membrane trafficking can destabilize CTL2 inclusions during mid to late infection stages ^51,54^. Hence, it appeared plausible that a potential instability of CpoS-deficient inclusions could stem from the mutant’s reduced ability to acquire sphingolipids. Its heightened susceptibility to SPT inhibition could then signify further aggravation of destabilization, which, if occurring early, could potentially result in infection clearance rather than cell death, thereby explaining our above-mentioned experimental observations. However, our past studies, based on live cell imaging, have suggested that CpoS-deficient inclusions only rupture at the onset of cell death ^55^, leaving unclear if CpoS-deficiency indeed causes membrane damage that can later cumulate into inclusion rupture triggering cell death or if observed ruptures were merely a consequence of cell death.

Given the lack of tools enabling detection of minor forms of inclusion membrane damage in *Chlamydia* spp., we once more employed the power of the recently established molecular genetic toolbox for *Chlamydia* to develop an innovative fluorescence microscopic tool capable of visualizing compromised inclusion membranes and bacteria released from damaged inclusions. This tool employs the split-GFP principle, where the N-terminal part of GFP (GFP1-10) and the C-terminal part (GFP11) are expressed separately but can associate to form a functional fluorescent protein upon co-localization. We generated CTL2 and CTL2-*cpoS*::*cat* derivatives expressing distinct variants of the Inc protein IncB or the *Chlamydia* outer membrane protein OmpA tagged with both FLAG and GFP11 (Fig 5A). In IncB-GFP11c, we placed multiple copies of GFP11 at IncB’s C-terminus, which is exposed to the host cell cytosol during infection. In IncB-GFP11int, the GFP11 copies were placed between IncB’s two transmembrane domains, a region presumed to face the inclusion lumen, while in OmpA-GFP11int, they were inserted into the predicted surface-exposed loop region between OmpA’s β-strands 11 and 12. Hence, during infection, the GFP11 tags of the constructs are expected to be displayed at the inclusion membrane, facing either towards the host cell cytosol (IncB-GFP11c) or the inclusion lumen (IncB-GFP11int), or at the surface of the bacteria (OmpA-GFP11int). These constructs can thus serve as positive control for the split-GFP system (IncB-GFP11c) or facilitate the detection of damaged inclusion membranes (IncB-GFP11int) or bacteria exposed to the host cell cytosol (OmpA-GFP11int) (Fig 5B).

**Figure 5.**
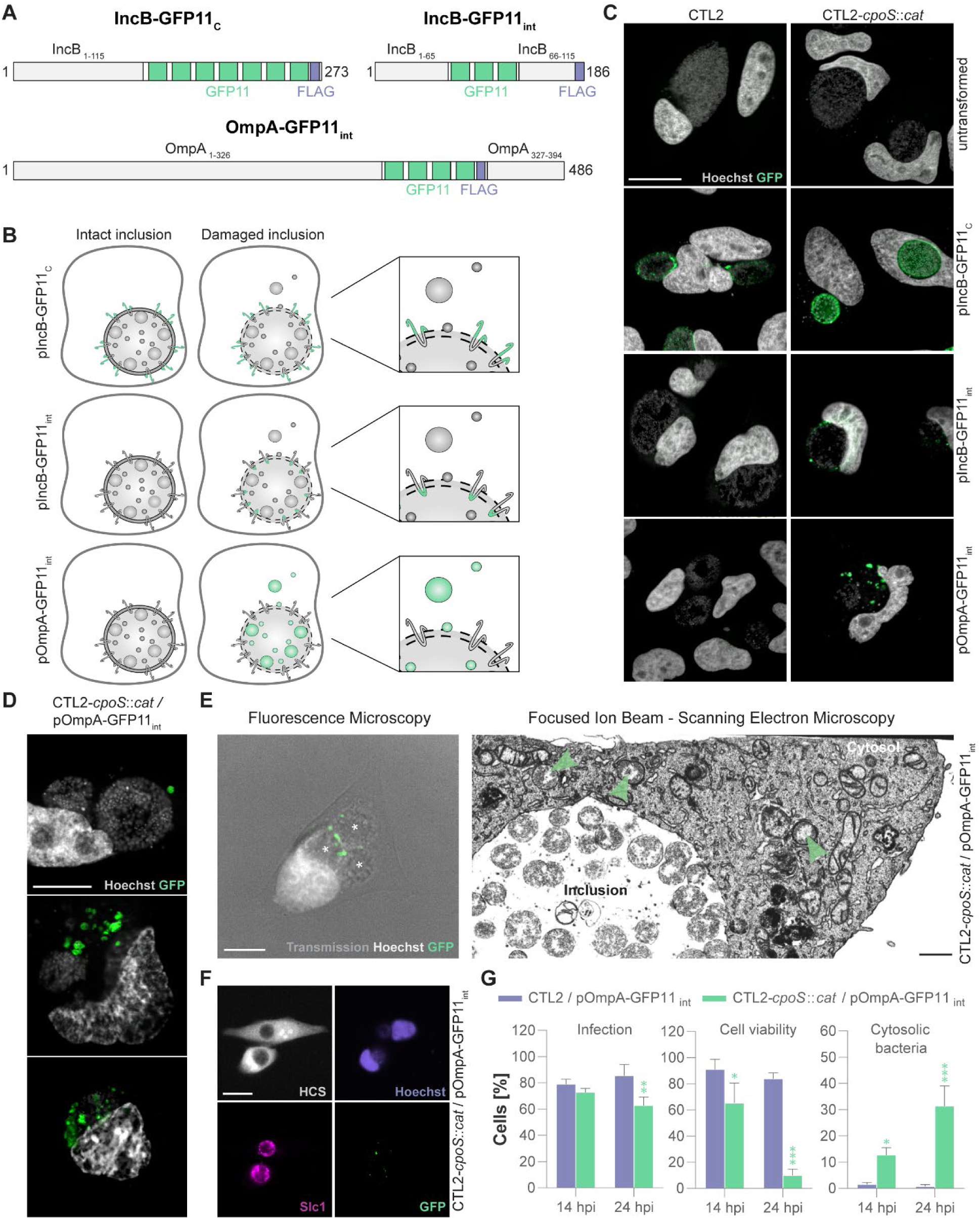
A novel reporter for inclusion damage revealed instability of CpoS-deficient inclusions. **(A)** Composition of the three GFP11-tagged constructs. **(B)** The principle of detecting inclusion damage using the split-GFP approach. **(C)** Fluorescence microscopic detection of inclusion damage during infection with CTL2-*cpoS*::*cat*. HeLa cells were transfected with a plasmid driving GFP1-10 expression, infected with the indicated strains (5 IFU/cell), and then fixed and imaged at 26 hpi (scale = 20 µm). **(D)** Confocal fluorescence microscopic images displaying different grades of inclusion damage. GFP1-10-expressing HeLa cells were infected with the indicated strain (10 IFU/cell) and then fixed and imaged at 24 hpi (scale = 10 µm). **(E)** FIB-SEM analysis validating inclusion damage at the ultrastructural level. GFP1-10-expressing HeLa cells were infected with the indicated strain and fixed at 24 hpi. A cell containing green-fluorescent bacteria was identified by fluorescence microscopy (left, scale = 10 µm, asterisks highlight inclusions) and subjected to FIB-SEM analysis (right, scale = 1 µm, one selected slice of the volume shown in Movie S1, green arrows highlight bacteria in the host cell cytosol). **(F-G)** Quantitative analysis of inclusion damage. GFP10-expressing HeLa cells were infected with the indicated strains (10 IFU/cell) and then fixed, stained (HCS, cells; Hoechst, DNA; Slc1, bacteria), and imaged at the indicated time points. Images were taken at a high-content imaging platform, which, despite low magnification, enabled sensitive detection of cells, nuclei, inclusions, and cytosolic bacteria. (F) Example image (scale = 25 µm). (G) The percentage of infected cells, surviving cells, and infected cells containing cytosolic bacteria was determined by manual image analysis (mean±SD, n=3, at least 100 cells per condition and replicate counted, 2-way ANOVA with Sidak’s, for each time point, indicated are significant differences compared to CTL2/pOmpA-GFP11int).

In infected HeLa cells, all three constructs were expressed and localized correctly to either the inclusion membrane or the bacteria (Fig S5A). In cells expressing cytosolic GFP1-10, IncB-GFP11c at the inclusion membrane of CTL2 became green fluorescent (Fig S5B). In contrast, no green fluorescence was observed in cells infected with CTL2 expressing IncB-GFP11int or OmpA-GFP11int (Fig S5B), consistent with the barrier function of the inclusion membrane. Critically, upon treating cells with membrane-permeabilizing agents, such as digitonin or high concentrations of DMSO, green-fluorescent spots emerged at IncB-GFP11int-positive inclusion membranes (Fig S5C). Moreover, inclusions of OmpA-GFP11int-expressing bacteria displayed green fluorescence post-treatment (Fig S5C). This fluorescence was faint and diffuse, implying a rupture of both the inclusion and bacterial membranes. With shorter treatment durations, individual green-fluorescent bacteria could still be discerned within some inclusions (Fig S5D).

Intriguingly, green-fluorescent spots, indicative of membrane damage, were also observed at CpoS-deficient IncB-GFP11int-positive membranes in the absence of permeabilizing agents (Fig 5C). Moreover, a subset of cells infected with OmpA-GFP11int-expressing CTL2-*cpoS*::*cat* harbored green-fluorescent bacteria (Fig 5C), often adjacent to seemingly intact non-fluorescent inclusions, likely representing early signs of inclusion damage (Fig 5D). Inclusion rupture, resulting in the majority of the bacteria exhibiting green fluorescence, was observed occasionally, but typically only in cells that appeared to be in the process of dying (Fig 5D). The leakage of CTL2-*cpoS*::*cat* into the host cell cytosol could further be confirmed at the ultrastructural level, when we selected cells with green-fluorescent bacteria for focused ion beam scanning electron microscopy (FIB-SEM) (Fig 5E). Moreover, a quantitative analysis by fluorescence microscopy revealed that at 14 hpi, coinciding with the onset of cell death in cultures infected with CTL2-*cpoS*::*cat*, and at 24 hpi, when extensive cell death was observed, about 12.7% and 25.3% of the infected cells, respectively, contained cytosolic bacteria (Fig 5F-G). Critically, this analysis also confirmed the near absence of cytosolic bacteria at these times during infection with CTL2.

In summary, our innovative microscopic tool, which enables the detection of early signs of inclusion damage and cytosolic *Chlamydia*, provided evidence that inclusions formed by CTL2 are usually stable and that CpoS indeed plays a key role in preserving inclusion integrity.

### SPT inhibition and replenishment of sphingoid bases modulated inclusion stability

After having confirmed that CpoS-deficient inclusions are unstable, we sought to investigate whether SPT inhibition indeed exacerbates inclusion damage. We infected GFP1-10-expressing HeLa cells with OmpA-GFP11int-expressing CTL2-*cpoS*::*cat* and parallelly treated them with MYR or LCS. In line with our previous findings, we observed robust protection from cell death and a substantial decrease in the proportion of cells containing inclusions at 24 hpi (Fig 6A). Unexpectedly, when observed at this time point, the percentage of infected cells containing cytosolic bacteria appeared unaffected by the presence of SPT inhibitors (Fig 6A). However, when we quantified inclusion damage at an earlier time point, at 18 hpi, we observed a strong inclusion-destabilizing effect due to SPT inhibition, with up to 70% of the treated infected cells harboring cytosolic bacteria. Collectively, these findings confirmed that SPT inhibition aggravates the instability of CpoS-deficient inclusions, with a majority of the inclusions becoming destabilized early on, leading to infection clearance rather than host cell death.

**Figure 6.**
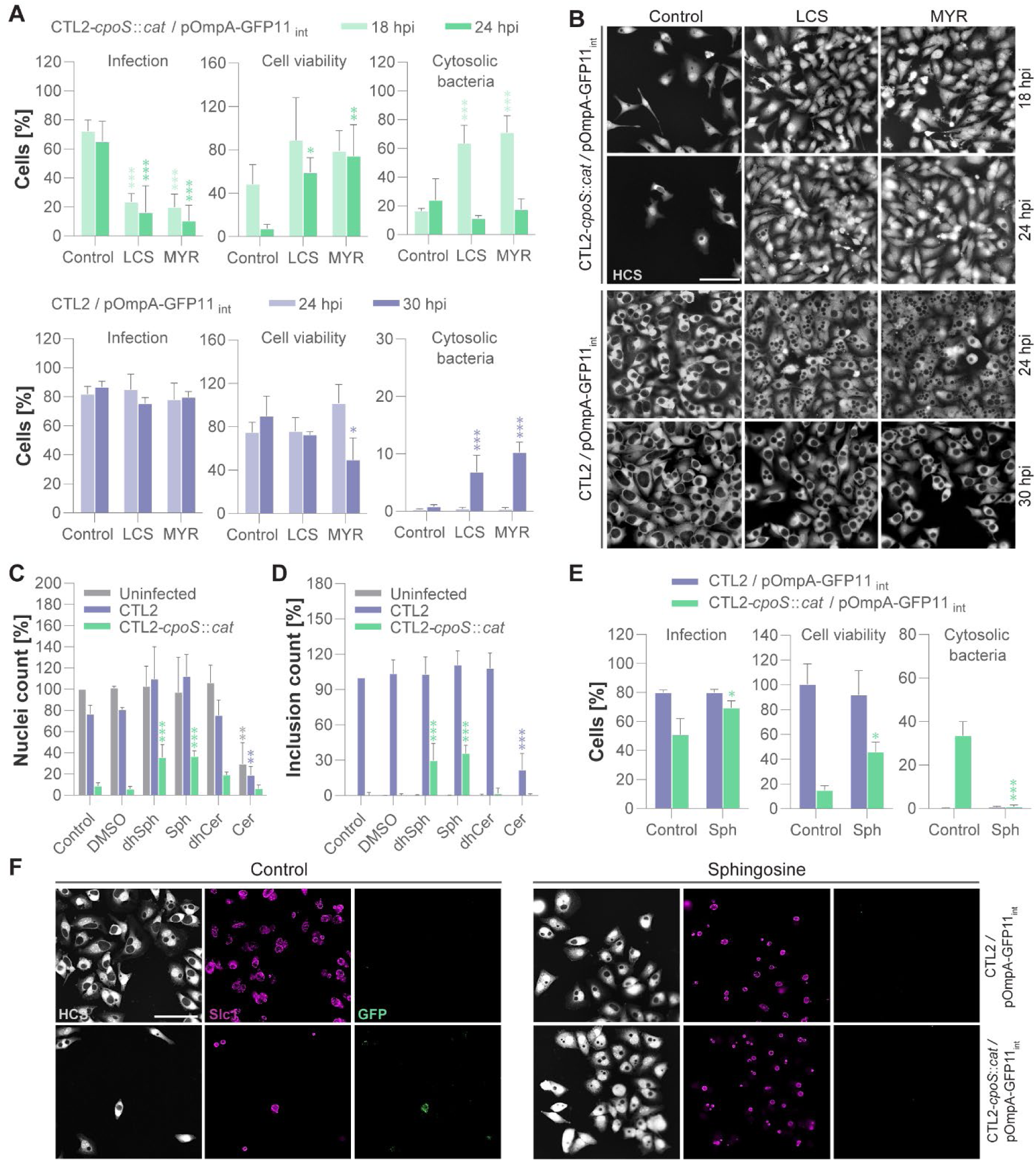
SPT inhibition and replenishment of sphingoid bases modulated inclusion stability. (A-B) Destabilizing effect of SPT inhibition on inclusions. GFP10-expressing HeLa cells were infected with the indicated strains (5 IFU/cell) and parallelly treated with MYR (10 µM) or LCS (125 µM) or left untreated (control). Cells were fixed, stained, and imaged at the indicated time points. (A) The percentage of infected cells, surviving cells, and infected cells containing cytosolic bacteria was determined by manual image analysis (mean±SD, n=3, at least 100 cells per condition and replicate counted, 2-way ANOVA with Sidak’s; for each time point, indicated are significant differences compared to the untreated control). (B) Representative images (scale = 100 µm). **(C-D)** Supplementation of growth media with sphingoid bases protected cells from *cpoS* mutant-induced death and restored bacterial growth. HeLa cells were infected with the indicated strains (4 IFU/cell), and parallelly treated with 5 µM of the indicated metabolites (dhSph, dihydrosphingosine; Sph, sphingosine; dhCer, dihydroceramide; Cer, ceramide; DMSO, solvent only) or left untreated (control). Nuclei counts (C) at 25.5 hpi are displayed normalized to the uninfected untreated control, while inclusion counts (D) are displayed normalized to the CTL2-infected untreated control (mean±SD, n=3, one-way ANOVA with Dunnett’s; for each infection condition, indicated are significant differences compared to the untreated control). **(E-F)** Stabilizing effect of sphingoid bases on CpoS-deficient inclusions. GFP10-expressing HeLa cells were infected with the indicated strains (5 IFU/cell) and parallelly treated with 5 µM sphingosine (Sph) or left untreated (control). Cells were fixed, stained (HCS, cells; Hoechst, DNA; Slc1, bacteria), and imaged at 24 hpi. (E) The percentage of infected cells, surviving cells, and infected cells containing cytosolic bacteria was determined by manual image analysis (mean±SD, n=3, at least 100 cells per condition and replicate counted, 2-way ANOVA with Sidak’s; for each strain, indicated are significant differences compared to the untreated control). (F) Representative images (scale = 100 µm).

When GFP1-10-expressing HeLa cells were infected with CpoS-proficient OmpA-GFP11int-expressing CTL2 and parallelly treated with MYR or LCS, we observed inclusion destabilization only at later stages of infection. Indeed, at 24 hpi, cytosolic bacteria were undetectable (Fig 6A), even though we observed an inclusion fusion defect in inhibitor-treated cells, as indicated by a high proportion of cells containing multiple inclusions (Fig 6B). At 30 hpi, cytosolic bacteria were detectable in a minor fraction of cells treated with SPT inhibitors, which in cells treated with MYR also coincided with a slight increase in host cell death (Fig 6A). Overall, this implies that CpoS-proficient inclusions exhibit greater resistance to the destabilizing effects of SPT inhibition. Moreover, destabilization at later stages seems to be linked to cell death, partially mirroring the phenotype of CpoS deficiency.

To confirm that the observed destabilization of CpoS-deficient inclusions resulted from diminished sphingolipid acquisition by inclusions, we provided cells with an excess of sphingolipid metabolites. The addition of C6-D-erythro-ceramide to the growth medium proved highly toxic to both infected and uninfected cells (Fig 6C), aligning with the pro-death signaling function of ceramides ^47^. Conversely, the addition of D-erythro-sphingosine or D-erythro-dihydrosphingosine protected cells from *cpoS* mutant-induced cell death and restored the growth of the mutant to a significant extent (Fig 6D). Notably, the addition of these sphingoid bases also stabilized CpoS-deficient inclusions (Fig 6E-F).

Together, these findings underscore the significance of sphingolipids in preserving inclusion stability. Critically, they also demonstrate that destabilization of inclusions at early infection stages leads to infection clearance while maintaining the viability of the host cell (Fig 7), a finding that could guide the development of future therapeutic approaches.

**Figure 7.**
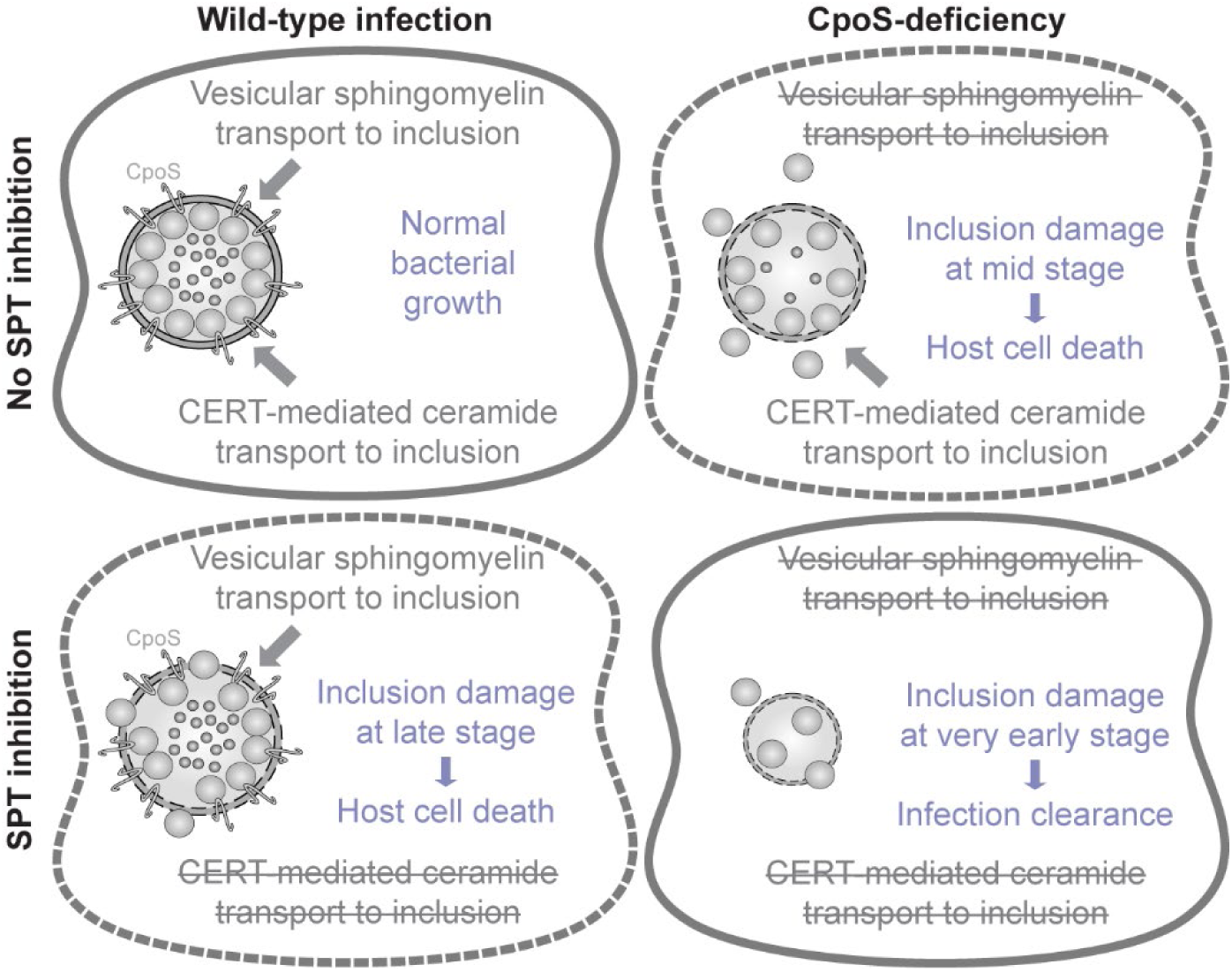
Model describing the role of CpoS in maintaining inclusion stability. *C. trachomatis* can acquire sphingolipids in two ways, by recruiting the host ceramide transfer protein CERT to acquire ceramides then converted to complex sphingolipids at the inclusion ^51^, and by hijacking sphingolipid-laden membrane vesicles and directing them to the inclusion ^52^. When CpoS is absent, the vesicular transport of sphingomyelin (and possibly other lipids) to the inclusion is compromised ^11^, leading to inclusion damage at mid stages of infection, followed by premature host cell death. We hypothesize that under conditions of SPT inhibition, the CERT-mediated transport is compromised, while the bacteria may still be able utilize the available lipid pool through CpoS-mediated modulation of host vesicular transport, leading to more minor inclusion damage that manifests at late stages of infection. A combination of SPT inhibition and CpoS-deficiency compromises both transport routes, resulting in early inclusion destabilization and clearance of infection while maintaining host cell viability. Hence, an effective therapeutic targeting of inclusion stability may have to aim for blocking both transport routes simultaneously.

## DISCUSSION

This study was motivated by our aim to decipher the molecular basis of the premature host cell death defense program suppressed by the *C. trachomatis* effector CpoS. Our journey began with an unbiased genetic screening approach, which led us down unforeseen paths, yet significantly advanced our molecular understanding of *Chlamydia*-host interactions, particularly the roles of CpoS and the inclusion in the evasion of host cell-autonomous immunity. Specifically, we (a) identified host genes with key functions during infection with *C. trachomatis* L2, also highlighting the role of HSPGs (Fig 1-2, Tables S2-S3), (b) uncovered host genes whose disruption conferred specific protection against the *cpoS* mutant (Fig 1-2, Tables S2-S3), (c) revealed presumably cytotoxic alterations in the levels and ratios of sphingolipid metabolites during infection with the mutant (Fig 3 and S3), (d) uncovered the mutant’s heightened dependence on host *de novo* ceramide synthesis (Fig 4), (e) confirmed the unstable nature of CpoS-deficient inclusions (Fig 5), and (f) established a role for CpoS in preserving inclusion stability via sphingolipid acquisition (Fig 6), thereby also highlighting possible targets for a potential therapeutic vacuole destabilization.

The identification of host genes with broad roles during CTL2 infection was not an explicit focus of this study but carried out primarily to allow excluding these genes from our subsequent studies in which we wanted to focus on genes contributing to the early cytotoxicity observed during CpoS-deficiency only. Nevertheless, our findings from this part of the study strongly highlighted the importance of HSPGs (Fig 1-2, Tables S2-S3), in accordance with previous work demonstrating a role for these cell surface molecules in facilitating the attachment and entry of CTL2 ^56–58^. Interestingly, previous CRISPR screens aimed at identifying host genes involved in various intracellular infections reported a similar trend with hits being heavily dominated by genes participating in pathogen attachment and invasion, rather than intracellular host-pathogen interactions (*^e.g.,^* ^23,26,59^). This possibly reflects a need for strong infection-inhibitory effects for robust enrichment in pooled screens ^60^. Nevertheless, some of our “general infection hits” may also have HSPG-independent roles during *C. trachomatis* infection. It is noteworthy that strains of *C. trachomatis* serovar E reportedly infect host cells in an HSPG-independent manner ^57^. Hence, future research could leverage this infection model to further clarify the infection-related roles of the identified genes. In this way, the findings of the genetic screen can also serve as a resource informing future experimental forays into *Chlamydia*-host interactions.

When it comes to the defense response in focus of our interest, we were surprised to find that the deficiencies conferring specific protection against the *cpoS* mutant did not affect known functions in pathogen recognition, immune signaling, or regulated forms of cell death (Fig 1-2, Tables S2-S3). However, considering CpoS’ function as a manipulator of membrane trafficking and mediator of lipid transport to the inclusion ^9,11,61,62^, we considered hits involved in trafficking and lipid synthesis also of special interest. As it relates to trafficking, TBC1D13, a GTPase-activating protein for RAB35 ^45^, may be particularly noteworthy. We recently reported that RAB35 is recruited to the CTL2 inclusion in a CpoS-dependent manner and plays a role in the CpoS-mediated dampening of the STING-dependent type I IFN response ^11^. Given that STING partially contributes to the cell death mediated by the *cpoS* mutant ^9^, it could be speculated that the protective effect of TBC1D13 deficiency might be due to it resulting in generally enhanced levels of active (GTP-bound) RAB35. This might compensate for the effects of CpoS-deficiency and dampen STING activation in infected cells. Yet, oddly, the immune sensor STING itself was not uncovered as a screening hit, suggesting that protection conferred by its deficiency is not potent enough to provide sufficient positive selection.

Screening hits in the *de novo* sphingolipid synthesis pathway specifically caught our attention. They were clearly specific to infections with the *cpoS* mutant (Fig 1-2, Tables S2-S3), and demonstrated robustness, as our hits included genes mediating two sequential enzymatic steps, catalyzed by SPT and KDSR, and genes encoding each molecular component of SPT (SPTSSA, SPTLC1, and SPTLC2) (Fig 3A). Alternative variants of SPT, including components such as SPTSSB and SPTLC3, do exist ^63^, yet are either not or only weakly expressed in HeLa cells according to the Human Protein Atlas ^64^. Moreover, there is genetic redundancy in subsequent steps of the *de novo* synthesis pathway and other ceramide-generating metabolic processes ^46^, which could explain why enzymes such as ceramide synthases and sphingomyelinases were not uncovered in the screen. Critically, a link between ceramide synthesis and host cell death seemed plausible, given the well-documented pro-apoptotic functions of ceramides ^47^.

Our initial validation experiments supported a scenario where CpoS-deficiency causes cell death through an accumulation of ceramides. Indeed, several inhibitors of ceramide-generating processes offered protection against *cpoS* mutant-induced cell death (Fig 3B-D). While our mass spectrometric analyses did not confirm the suspected metabolite accumulation at time points preceding or coinciding with the onset of cell death (Fig 3E and S3), they revealed an increase in the dihydroceramide/ceramide ratio (Fig 3E), another metabolic alteration known to cause cell death ^48^. A contribution of this alteration to the *cpoS* mutant-triggered premature host cell death seems plausible, given our finding that SKI-II, which increases the dihydroceramide/ceramide ratio by inhibiting dihydroceramide desaturase ^49^, induced death in cells infected with CTL2 (Fig 3B-C). Similar effects on host cell viability were not seen after exogenous supplementation of C6-D-erythro-dihydroceramide (Fig 6C). Yet, this finding does not rule out a possible role of endogenous dihydroceramide species. In addition to these considerations, our findings with GW4869, which blocked *cpoS* mutant-induced cell death without causing notable defects in bacterial growth (Fig 3B-D and 4A-C), and the observed accumulation of S1P (Fig 3E and S3), a metabolite typically produced from ceramides derived from sphingomyelinase activity, leave open the possibility for a contribution of sphingomyelinases in *cpoS* mutant-induced cell death.

Beside the observation of these potentially cytotoxic metabolic alterations, which may have been ameliorated by the pharmacologic inhibitors used, our further analyses demonstrated that SPT inhibition also curbed the growth of the *cpoS* mutant by aggravating inclusion instability at early infection stages (Fig 6A). While CpoS deficiency had previously been suggested to cause premature inclusion rupture ^10^, the molecular basis of this putative inclusion membrane damage remained unknown. Furthermore, given that our prior findings suggested rupture to occur only concurrently with cell death ^55^, it remained unclear if CpoS-deficient inclusions were indeed inherently unstable, as the observed ruptures could have also resulted as a byproduct of host cell death. Earlier research on vacuolar instability in *Chlamydia* primarily relied on the fluorescence microscopic detection of dispersed intracellular bacteria and/or discontinuous inclusion membrane staining ^10,51,54,65–69^. Other studies used the influx of GFP expressed in the host cell cytosol as an indicator of membrane rupture ^5,70^. However, a system for the specific and sensitive detection of early signs of inclusion instability was lacking, prompting us to develop an appropriate tool. We opted to develop the split-GFP-based reporter, as this innovative approach offers versatility in detecting various signs of damage, including damaged inclusion membranes and individual bacteria released from inclusions. The tool enables quantitative analyses of inclusion damage and should be compatible with live cell imaging as well. Overall, we predict that this novel tool will be an important addition to the available toolbox for *Chlamydia* spp., as by providing an ability to monitor inclusion stability and the host response to bacteria entering the host cell cytosol, it will also revolutionize our ability to mechanistically dissect the host cell-autonomous defense against *Chlamydia* infection. Crucially, the tool already filled an important knowledge gap by providing experimental evidence that CpoS-deficient inclusions are indeed unstable, and by offering insights into the peculiar nature of this inclusion damage, initially characterized by the release of individual bacteria rather than an immediate rupture of the vacuole (Fig 5C-G).

Earlier studies suggested that blockades in membrane trafficking or ceramide synthesis can cause some degree of inclusion destabilization ^51,54^. This led us hypothesize that the previously discovered reduced capacity of CpoS-deficient inclusions to acquire sphingolipids ^11^ could be the base for their instability. Indeed, we found that SPT inhibition further destabilized CpoS-deficient inclusions, while supplementing media with sphingoid bases, specifically sphingosine and possibly also dihydrosphingosine, stabilized them, thereby restoring to some extent host cell viability and bacterial growth (Fig 6C-F). Although SPT inhibition also destabilized CTL2 inclusions, this only occurred at later infection stages (Fig 6A). Critically, inclusion destabilization during mid-stages of infection with the *cpoS* mutant or at late stages in SPT-inhibited CTL2-infected cells led to host cell death (Fig 5G and 6A). Conversely, inclusion destabilization at early stages, in SPT-inhibited *cpoS* mutant-infected cells, led to infection clearance while preserving the host cells (Fig 6A). This finding may in part also explain why CERT-deficiency in the CRISPR screen appeared to have a tendency for protecting cells from *cpoS* mutant-induced cell death, while making them more susceptible to CTL2-induced cytotoxicity (Fig S4A). If the reduced cell death observed in SPT-inhibited *cpoS* mutant-infected cells may in part be due to the inhibitor’s effects on metabolite levels remains to be deciphered, as do the molecular details of infection clearance. Nevertheless, our findings suggest that strategies destabilizing inclusions at early infection stages could effectively eradicate the bacteria without harming the host cells (Fig 7).

There are several important points future research will need to address. For instance, it remains to be clarified if the observed alterations in sphingolipid metabolites in cells infected with the *cpoS* mutant are a consequence of the altered interactions of the bacteria with the host cellular lipid pool, or if they could possibly reflect the action of a host defense response triggered by the presence of the bacteria in the cytosol. In the latter case, one may assume perturbations in sphingolipid metabolites to potentially have a wider role in the cell-autonomous defense against intracellular pathogens. On the other hand, considering that an increased dihydroceramide/ceramide ratio in other instances was found to cause cell death by disturbing the integrity of autolysosomal membranes ^48^, it also needs to be determined if such metabolic alteration could directly contribute to the destabilization of CpoS-deficient inclusion membranes.

In conclusion, this study has substantially advanced our understanding of how *C. trachomatis* circumvents host cell-autonomous immunity. It underscores the importance of the inclusion as an intracellular sanctuary and the role of CpoS in preserving this refuge by ensuring an adequate supply of sphingolipids. The finding that early inclusion destabilization can eradicate infection without harming the host cells may pave the way for innovative therapeutic strategies. In this regard, the split-GFP tool for assessing inclusion damage is an invaluable addition to the existing toolbox for *Chlamydia*, as its future applications will aid in further deepening our molecular understanding of inclusion stability and in developing therapeutic means for inclusion destabilization.

## MATERIALS AND METHODS

### Cell culture

HeLa (ATCC CCL-2), Vero (ATCC CCL-81), and HEK293T (ATCC CRL-3216) cells were cultivated in Dulbecco’s Modified Eagle’s Medium (Gibco), supplemented with 10% heat-inactivated fetal bovine serum (Gibco). In selected experiments, including those involving an analysis of cell death via the resazurin assay, a DMEM medium devoid of phenol red was used to avoid interference with measurements of fluorescence. Cultures were maintained in a humidified incubator (37°C, 5% CO2) and routinely tested for contamination with *Mycoplasma* spp. using a commercial PCR-based *Mycoplasma* detection assay (VenorGeM, Minerva).

### Treatment with pharmacological inhibitors or metabolites

The following inhibitors targeting enzymes in ceramide synthesis, metabolism, or transport were used: fumonisin B1 (Cayman), NVP-231 (Cayman), L-cycloserine (Cayman), myriocin (Cayman), D-erythro-MAPP (Cayman), GW4869 (Cayman), SKI-II (BioVision), desipramine hydrochloride (BioVision), (1R,3S)-HPA-12 (Cayman), and golgicide A (Cayman). Additionally, we used the following sphingolipid metabolites: D-erythro-sphingosine (Merck), D-erythro-dihydrosphingosine (Merck), C6-D-erythro-dihydroceramide (Merck), C6-D-erythro-ceramide (Merck). Stock solutions of inhibitors and metabolites were prepared in DMSO, except for fumonisin B1 and D-erythro-MAPP, which were dissolved in ethanol. The inhibitors and metabolites were added to cell cultures at concentrations and for durations as specified in the respective figure legends.

### *Chlamydia* strains and general infection procedure

Infection experiments were carried out with *C. trachomatis* L2/434/Bu (ATCC VR-902B), including the wild-type strain (CTL2), a strain carrying an insertional disruption of *cpoS* (CTL2-*cpoS*::*cat*), and further genetic derivatives as listed in Table S4. Two different procedures were used to prepare infection inocula, including crude preparations (used in infection experiments involving the split-GFP system or GFP-expressing bacteria) and highly pure, density-gradient purified EB preparations (used in all other experiments, including the screen). These procedures have been described in detail in a previous study ^11^. Bacteria obtained through either of the two methods were resuspended in SPG (sucrose-phosphate-glutamate) buffer (75 g/l sucrose, 0.5 g/l KH2PO4, 1.2 g/l Na2HPO4, 0.72 g/l glutamic acid, pH 7.5) and stored at -80°C. Furthermore, the bacterial preparations were titered (as previously described ^11^) and confirmed to be free from *Mycoplasma* contamination (as described above for cell lines). To conduct infections, cells were typically seeded in multi-well plates, followed by the addition of bacteria (number of inclusion forming units (IFUs)/cell as specified), centrifugation (1500 x g, 30 min, 23°C), and incubation (37°C, 5%CO2) for the indicated periods of time. The centrifugation step was omitted for infections conducted in MatTek dishes. Moreover, infections conducted in the context of the CRISPR screen were done in suspension, as described below.

### CRISPR screen

#### Generation of Cas9-expressing cells

HeLa cells were lentivirally transduced with pLenti-Cas9-T2A-Blast-BFP (Addgene #196714) to express a codon optimized, wild-type SpCas9, flanked by two nuclear localization signals, and linked to a blasticidin-S-deaminase – mTagBFP fusion protein via a self-cleaving peptide. Following selection with blasticidin, a stable BFP+ population was isolated through repeated sorting for BFP expressors.

#### Guide library construction

The genome-wide Brunello sgRNA library ^71^ (which targets 19,114 genes and comprises a total of 77,441 sgRNAs, including about four sgRNAs per gene and 1000 non-targeting control sgRNAs) was synthesized as 79 bp long oligos (CustomArray, Genscript). The oligo pool was double-stranded by PCR to include an A-U flip in the tracrRNA ^72^, 10 nucleotide long, untranscribed random sequence labels (RSLs), and an i7 sequencing primer binding site ^73^. The PCR product with the sequence ggctttatatat**cttgtggaaaggacgaaacaccgNNNNNNNNNNNNNNNNNNNNgtttaagagctagaaatagca agtttaaataaggct**agtccgttatcaacttgaaaaagtggcaccgagtcggtgctttttt<u>GATCGGAAGAGCACACGTCTGAACTCCAGTCAC</u>NNNNNNNNNNaagcttggcgtaactagatcttgagacaaa (array oligo in bold, i7 primer binding underlined) was cloned by Gibson assembly into pLenti-Puro-AU-flip-3xBsmBI ^73^ (Addgene #196709). The plasmid library was sequenced to confirm representation and packaged into lentivirus in HEK293T cells using plasmids psPAX2 (Addgene #12260, gift from Didier Trono) and pCMV-VSV-G (Addgene #8454, gift from Bob Weinberg). The virus-containing supernatant was concentrated around 40-fold with Lenti-X concentrator (Takara), aliquoted, and stored in liquid nitrogen.

#### Library virus titration and large-scale transduction

The functional titer of the library virus was estimated from the fraction of surviving cells after transduction of target cells with different amounts of virus and puromycin selection. For the screen, Cas9-BFP-expressing target cells were transduced in duplicate with the library virus at an approximate multiplicity of infection of 0.3 and a coverage of 1,000 cells per guide in the presence of 2 µg/ml polybrene. Transduced cells were selected with 2 µg/ml puromycin from day 2 to day 4 post transduction, and then frozen at day 6 post transduction and stored at -150°C. Throughout this procedure, the cell count was kept at ≥ 80 million per replicate to ensure full library coverage.

#### Selection of cells resistant to infection-induced cell death

Transduced cells were thawed and transferred to large culture flasks for incubation at 37°C, 5%CO2. A cell count conducted 12 hours post-thawing revealed over 90 million live cells. When the cultures approached confluency (4-5 days post-thawing), cells from all flasks were harvested, pooled, and counted. A total of 1×10^8^ cells were transferred to a 50 ml tube, washed once with Dulbecco’s phosphate-buffered saline (DPBS), pelleted (800 × g, 10 min), and stored at -80°C (= pre-selection cell population). The remaining cells were divided into four batches (each 1.14×10^8^ cells) to establish the four sample groups: UI30h, UI60h, KO30h, WT60h. The groups KO30h and WT60h were then infected by addition of 100 µl SPG buffer containing 3.41×10^9^ IFUs (30 IFU/cell) of CTL2-*cpoS*::*cat* or CTL2, respectively, while the groups UI30h and UI60h were treated with 100 µl bacteria-free SPG. To enhance infection, the cell suspensions were subjected to two rounds of mixing (by pipetting) and centrifugation (1500 x g, 10 min, 23°C). Subsequently, the cells were transferred to large culture flasks and incubated at 37°C, 5%CO2. At the time of cell collection, *i.e.*, at 30 hpi (UI30h and KO30h) or 60 hpi (UI60h and WT60h), the surviving cells were washed twice with DPBS in the flasks (to remove dead cells and cell debris), harvested, counted, washed once with DPBS, pelleted (800 x g, 10 min), and stored at -80°C (= selected cell populations). Two independent biological replicates (R1+R2) of this procedure were conducted, starting from the two independent lentiviral transductions.

#### Isolation of genomic DNA, library preparation, and sgRNA sequencing

Genomic DNA was isolated from all cell pellets using the QIAamp DNA Blood Maxi Kit (Qiagen). Prior to following the kit’s instructions, RNase A (800 µg/10 million cells) was added to all samples. DNA was quantified using Qubit dsDNA BR Assay Kit (Thermo-Fisher-Scientific). Guide and UMI sequences were amplified by PCR, as previously described ^73^, using modified primers PCR2_FW (5’-ACACTCTTTCCCTACACGACGCTCTTCCGATCTCTTGTGGAAAGGACGAAACAC-3’) and PCR3_fw (5’-AATGATACGGCGACCACCGAGATCTACAC**[i5]**ACACTCTTTCCCTACACGACGCTCT-3’), respectively. The amplicon was sequenced on Illumina NovaSeq, reading 20 cycles Read 1 with custom primer 5’-CGATCTCTTGTGGAAAGGACGAAACACCG-3’, 10 cycles index read i7 to read the RSL, and six cycles index read i5 to read the sample barcode. The sequencing data were analyzed using the MAGeCK software (v.0.5.6) ^74^.

### Transient and stable expression of GFP1-10 in human cells

Plasmid pcDNA3.1-Cyto-GFP1-10 ^75^, enabling transient expression of cytosolic GFP1-10, was kindly provided by Kevin Hybiske (University of Washington). HeLa cells were transfected with this plasmid using jetPRIME transfection reagent (Polyplus) with a medium exchange after 2-3 hours. To enable stable expression of GFP1-10 in HeLa cells, a lentiviral expression system was used. The sequence encoding GFP1-10 was PCR-amplified from pcDNA3.1-Cyto-GFP1-10 using the primers GFP1-10_F (5’-TATGTCGACATGGTTTCGAAAGGCGAGGA-3’) and GFP1-10_R (5’-GGCCTCGAGTTATTTCTCGTTTGGGTCTT-3’). The PCR product was then digested with EcoRI and BamHI and ligated into EcoRI/BamHI-digested lentiviral vector pLVX-Puro (Takara). Subsequently, HEK293T cells were co-transfected with the resulting vector pLVX-Puro-GFP-1-10 and the packaging and envelope plasmids using Lenti-X Packaging Single Shots (Takara). Supernatants containing lentiviral particles were collected at 48 and 72 hours post-transfection, filtered (0.45 µm), and used to transduce HeLa cells in the presence of 4 µg/ml polybrene (Merck). At 24 hours post-transduction, the medium was replaced with a medium containing 1 µg/ml puromycin (Gibco) to select for cells that had been stably transduced. Cell clones were generated by limiting dilution and screened for robust GFP1-10 expression. This was based on the fluorescence microscopic detection of GFP signals following infection with strain CTL2/pIncB-GFP11c (described below).

### Generation of GFP11-expressing *Chlamydia* strains

Vector pTL2-tetO-IncB-GFP11x7-FLAG ^75^, enabling inducible expression of IncB (CT232) tagged at its C-terminus with seven copies of GFP11 (RDHMVLHEYVNAAGIT) and the FLAG epitope (DYKDDDDK), *i.e.*, construct IncB-GFP11c, was a kind gift from Kevin Hybiske (University of Washington). Vector pTL2-tetO-CTL0050-GFP11x4-FLAG-CTL0050, enabling inducible expression of OmpA (CTL0050) with four copies of GFP11 (RDHMVLHEYVNAAGIT) and a FLAG epitope (DYKDDDDK) inserted into OmpA’s predicted surface-exposed loop region between the β-strands 11-12, *i.e.*, construct OmpA-GFP11int, and vector pTL2-tetO-IncB-GFP11x3-IncB-FLAG, enabling inducible expression of IncB (CTL0484) with a C-terminal FLAG epitope and three copies of GFP11 inserted into IncB’s predicted inclusion lumen-exposed sequence, *i.e.*, construct IncB-GFP11int, were generated as follows: Gene blocks encoding the modified OmpA and IncB genes (Methods S1) were purchased (Thermo-Fisher Scientific). The genes were PCR-amplified (OmpA: 5’-TATAGGTATGCGGCCGCATGAA-3’, 5’-GCGGTAAGTGTCGACTTATTAG-3’; IncB: 5’-ATTATCGGCCGATGGT-3’, 5’-CGCGTGTCGACTTATTAC-3’), digested with NotI and SalI (OmpA) or EagI and SalI (IncB), and inserted into EagI/SalI-digested vector pTL2-tetO-IncB-GFP11x7-FLAG. The vectors encoding these three constructs were then transformed into CTL2 and CTL2-*cpoS*::*cat* using the CaCl2 approach ^76^. Transformed bacteria were selected in the presence of penicillin G (Merck) and plaque-purified ^9^. The presence of the correct plasmid in the clones was verified by PCR (pTL2-F: 5’-TCGATTTTTGTGATGCTCGTCAG-3’, pTL2-R1: 5’-ATATGATCTCTACCTCCAGAGCC-3’) and analysis of PCR products by agarose gel electrophoresis. In experiments using these strains, the expression of GFP11-tagged IncB or OmpA was induced by the addition of 4 ng/ml anhydrotetracycline (Clontech) at 0-2 hpi.

### Fluorometric quantification of host cell viability and bacterial growth

The ability of cells to convert non-fluorescent resazurin into fluorescent resorufin was used as an indicator of cell viability. In brief, resazurin (Merck; 0.15 mg/ml in DPBS) was added to the cells at a volumetric ratio of 1:5 at 24 hpi, followed by further incubation for 1.5 hours (37°C, 5% CO2). Resorufin fluorescence was then measured (excitation: 560 nm, emission: 590 nm, bandwidth: 10 nm) and normalized to the fluorescence detected in untreated cells to calculate cell viability as a percentage. To quantify intracellular bacterial growth fluorometrically, infections were conducted using rsGFP-expressing derivatives of CTL2 and CTL2-*cpoS*::*cat* (Table S4). At indicated times, the cells were fixed with 4% formaldehyde for 20 minutes at room temperature. Immediately afterwards, GFP fluorescence was measured (excitation: 490 nm, emission: 510 nm, bandwidth: 10 nm) and normalized to the fluorescence detected in untreated infected cells to calculate bacterial growth as a percentage. All assays described above were conducted in 96-well plates, and fluorescence measurements were made using a Spark plate reader (Tecan).

### Quantification of sphingolipids

For the quantification of sphingolipids, HeLa cells seeded in 6-well plates were infected with CTL2 or CTL2-*cpoS*::*cat* (5 IFU/cell) in the presence or absence of GW4869 (12.5 µM). At the indicated time points, cell extracts were prepared (from about 1×10^6^ cells per sample). In brief, cells were washed with ice-cold DPBS and lysed in ice-cold methanol (HPLC-grade, ≥99.9%, Sigma-Aldrich). Lysates were then collected with a cell scraper and stored at -80°C until further processing. Sphingolipids were extracted using 1.5 ml methanol/chloroform (1:1, v:v), containing the internal standards d7-dihydrosphingosine (d7-dhSph), d7-sphingosine (d7-Sph), d7-sphingosine 1-phosphate (d7-S1P), 17:0 ceramide (d18:1/17:0), d31-16:0 sphingomyelin (d18:1/16:0-d31), 17:0 glucosyl(β) ceramide (d18:1/17:0) and 17:0 lactosyl(β) ceramide (d18:1/17:0) (all from Avanti Polar Lipids) ^77^. Final extracts were subjected to sphingolipid quantification by liquid chromatography tandem-mass spectrometry (LC-MS/MS) applying the multiple reaction monitoring (MRM) approach. Chromatographic separation was achieved on a 1290 Infinity II HPLC (Agilent Technologies) equipped with a Poroshell 120 EC-C8 column (3.0 x 150 mm, 2.7 µm; Agilent Technologies) guarded by a pre-column (3.0 x 5 mm, 2.7 µm) of identical material. MS/MS analyses were carried out using a 6495C triple-quadrupole mass spectrometer (Agilent Technologies) operating in the positive electrospray ionization mode (ESI+). Chromatographic conditions and settings of the ESI source and MS/MS detector have been published elsewhere ^78^. The mass transitions used for analysis of sphingolipid subspecies are given in Table S5. Peak areas of Cer, dhCer, SM, dhSM, HexCer and LacCer subspecies, as determined with MassHunter software (Agilent Technologies), were normalized to those of their internal standards followed by external calibration. DhSph, Sph, and S1P were directly quantified via their deuterated internal standards. Quantification was performed with MassHunter Software (Agilent Technologies).

### Measurements of sphingomyelinase activity

Neutral sphingomyelinase activity in cell lysates was measured using a commercial colorimetric sphingomyelinase assay kit (Merck). In brief, HeLa cells were lysed at 18 hpi by incubation in sterile water for 20 min on ice. The lysates were then mixed with a concentrated buffer solution to achieve final concentrations of 25 mM Tris/HCl (pH 7.4) and 150 mM NaCl. This mixture was sonicated for 1 minute on ice and centrifuged at 17,000 x g at 4°C for 20 minutes to remove insoluble materials. Subsequently, sphingomyelinase activity was quantified as per the instructions provided in the kit. Moreover, the protein content was quantified using a BCA protein assay kit (Pierce), which enabled the calculation of activity as mU per mg protein.

### Fluorescence microscopy

For standard fluorescence microscopy evaluations (not coupled with electron microscopy), cells were cultivated in 96-well plates or on glass coverslips placed into 24-well plates and fixed for 20 minutes with 4% formaldehyde in DPBS. Subsequently, the cells were permeabilized for 15 minutes with 0.2% Triton X-100 in DPBS, incubated for 20 minutes in blocking solution (2-3% BSA in DPBS), and then for 1 hour in blocking solution containing primary antibodies (rabbit-anti-Slc1 (gift from Raphael Valdivia ^79^; 1:1000) or rabbit-anti-FLAG (Merck, F7425; 1:1000)). After washing thrice with DPBS, the cells were incubated for 1 hour in blocking solution containing the DNA dye Hoechst 33342 (Invitrogen; 2-10 µg/ml), Alexa Fluor 555-conjugated secondary antibodies (Invitrogen; 1:250-1:1000), and the whole cell dye HCS CellMask Deep Red Stain (Invitrogen; 5 ng/ml). Subsequently, the cells were washed thrice with DPBS again. Cells grown and stained on glass coverslips were transferred to microscope slides and embedded in ProLong Glass Antifade Mountant (Invitrogen). All incubation steps were conducted at room temperature. In some experiments, staining with the whole cell dye or with antibodies was omitted. Images were captured using various microscopy systems, including an epifluorescence microscope (Zeiss Axio Imager.Z2), a confocal microscope (Leica SP8), and a high-content imaging platform (Molecular Devices ImageXpress Micro XL system). Automatic processing of images recorded at the ImageXpress system and detection of chlamydial inclusions and host cell nuclei were performed with CellProfiler (version 4.0.7) ^80^. To quantify the extent of inclusion damage, images were analyzed manually. A minimum of 100 cells were analyzed per condition and biological replicate. Cells were classified as “uninfected” and “infected” based on the presence of inclusions. Within the group of “infected” cells, we further differentiated between cells that “contained cytosolic bacteria” and those that “did not contain cytosolic bacteria”.

### FIB-SEM imaging

GFP1-10-expressing HeLa cells were seeded in gridded 35 mm glass bottom MatTek dishes (MatTek, P35G-1.5-14-C-GRD) and subsequently transfected with pcDNA3.1-Cyto-GFP1-10, infected with the indicated strains, and treated with anhydrotetracycline, using above-described standard procedures. At 23.5 hpi, Hoechst 33342 (10 µg/ml) was added to the dishes, and at 24 hpi, the cells were fixed with 4% paraformaldehyde in 0.1 M PHEM buffer. Fluorescence microscopic analysis was conducted using a Leica DMi8 inverted Thunder widefield microscope, equipped with a LEICA DFC 9000 GTC camera and operated by Leica Application Suite X 3.8.2. Subsequently, the cells were postfixed with 2.5% glutaraldehyde in 0.1 M PHEM buffer. Samples were processed for FIB-SEM using a PELCO Biowave Pro+ microwave tissue processor (Ted Pella) following a previously described procedure ^81^, with minor modifications: calcium was not used during fixation and the contrasting step with lead aspartate was omitted to reduce the risk of overstaining. The samples were embedded in a thin disc of Durcupan resin, mounted on an SEM stub with epoxy and silver glue, and coated with 5 nm platinum. Before imaging, the area of interest was coated with a 1 µm thick protective layer of platinum. The volume was imaged using a FEI Scios DualBeam system, with the acquisition automated using the Auto Slice & View software (ver. 4.1.1.1582). Images were captured with the electron beam operating at 2 kV and 0.2 nA, detected with the T1 in-lens BSE detector. The voxel size (XxYxZ) of the volume was 9.6 x 9.6 x 20 nm. The volume was further registered and processed in ImageJ using the plugins “Linear Stack Alignment with SIFT” and “MultiStackReg”. After registration, the volume was converted into an MRC file and the header was modified to include pixel size information. All modeling was performed using the IMOD software package version 4.9.13 (Mastronade).

### Statistical analysis

Statistical analyses, other than those described for the CRISPR screen, were conducted in GraphPad Prism 8. The following statistical tests were used when indicated in the figure legends: one-way or two-way ANOVA with the indicated multiple comparison post hoc tests. The following significance levels were considered: *, p<0.05; **, p<0.01; *** p<0.001; ns, not significant.

## ACKNOWLEDGEMENTS AND FUNDING SOURCES

We wish to express our gratitude to Isabelle Dérre (University of Virginia), Kevin Hybiske (University of Washington), and Raphael Valdivia (Duke University) for their generosity in sharing reagents. The support by Daniel Herrmann in the LC-MS/MS analysis of sphingolipids is acknowledged. We also acknowledge the Umeå Centre for Electron Microscopy (UCEM), hosting the Umeå node of SciLifeLab Integrated Microscopy Technologies (IMT) at Umeå University, the Biochemical Imaging Center at Umeå University (BICU), and the Swedish National Microscopy Infrastructure (NMI, VR-RFI 2019-00217), for providing access to facilities and technical assistance with microscopy. Furthermore, we wish to acknowledge the Chemical Biology Consortium Sweden (CBCS) and its node at Umeå University for providing access to a high-content imaging platform. CBCS is a Swedish national research infrastructure funded by the Swedish Research Council (VR-2021-00179), SciLifeLab, and its hosting universities. The work performed at Umeå University was supported by Vetenskapsrådet (VR-2016-06598, VR-2018-02286, VR-2021-06602, VR-2022-00852), Kempestiftelserna (JCK-1834 and JCK-2031.2), and a Strategic Grant provided by Umeå University Medical Faculty. Part of this work was carried out at the SciLifeLab CRISPR Functional Genomics unit (CFG) at Karolinska Institutet, funded by SciLifeLab. CFG acknowledges support from the National Genomics Infrastructure, SNIC (project snic2017/7-265), and the Uppsala Multidisciplinary Center for Advanced Computational Science (UPPMAX). Furthermore, part of this work was funded by the Deutsche Forschungsgemeinschaft (DFG) – RTG 2581 “SphingoInf” (project number 417857878) granted to BK.

## AUTHOR CONTRIBUTIONS

Conceptualization, MRBS, LHJ, BSS; Investigation, MRBS, LHJ, MM, CLS, SD, FS, SH, NVGV, AC, PM, MÖ, SM, SFA, BSS; Formal analysis, MRBS, LHJ, MM, FS, MÖ, BS, BSS; Visualization, MRBS, LHJ, BSS; Writing – Original Draft, MRBS, LHJ, BSS; Writing – Review & Editing, MRBS, LHJ, FS, BS, BSS; Supervision, BS, BK, BSS; Funding acquisition, BK, BSS. All authors read and approved the final manuscript.

