## Supplemental Figures and Methods for "A genome-wide genetic screen identified targets for destabilizing the parasitophorous vacuole of *Chlamydia trachomatis*"

Short title: *Chlamydia* CpoS maintains inclusion integrity via sphingolipid acquisition

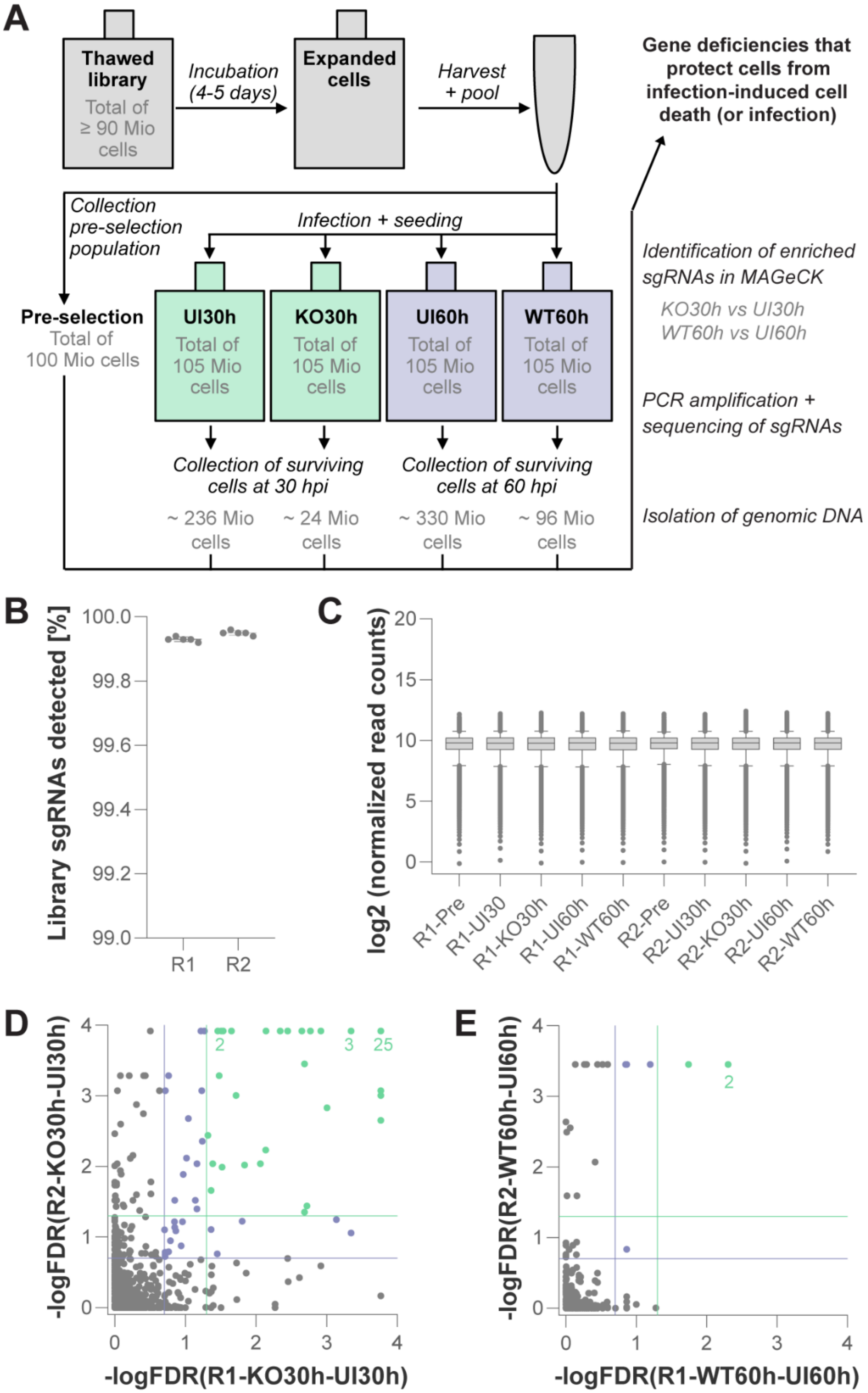

A
Thawed library
Total of ≥ 90 Mio cells
Incubation (4-5 days)
Expanded cells
Harvest + pool
Gene deficiencies that protect cells from infection-induced cell death (or infection)
Collection pre-selection population
Infection + seeding
Pre-selection
Total of 100 Mio cells
UI30h
Total of 105 Mio cells
KO30h
Total of 105 Mio cells
UI60h
Total of 105 Mio cells
WT60h
Total of 105 Mio cells
Identification of enriched sgRNAs in MAGeCK
KO30h vs UI30h
WT60h vs UI60h
PCR amplification + sequencing of sgRNAs
Collection of surviving cells at 30 hpi
Collection of surviving cells at 60 hpi
~ 236 Mio cells
~ 24 Mio cells
~ 330 Mio cells
~ 96 Mio cells
Isolation of genomic DNA
B
Library sgRNAs detected [%]
100.0
99.8
99.6
99.4
99.2
99.0
R1
R2
C
log2 (normalized read counts)
20
15
10
5
0
R1-Pre
R1-UI30
R1-KO30h
R1-UI60h
R1-WT60h
R2-Pre
R2-UI30h
R2-KO30h
R2-UI60h
R2-WT60h
D
-logFDR(R2-KO30h-UI30h)
-logFDR(R1-KO30h-UI30h)
2
3
25
E
-logFDR(R2-WT60h-UI60h)
-logFDR(R1-WT60h-UI60h)
2

**Figure S1. Selection procedure in the CRISPR screen and identification of gene deficiencies that protect human cells from *C. trachomatis*-induced cytotoxicity. (A)** Detailed schematic illustration of the selection procedure in the CRISPR screen (UI, uninfected; KO, knock-out (CTL2-*cpoS*::*cat*); WT, wildtype (CTL2)). **(B)** Proportion of library sgRNAs detected in the sequenced samples, including Pre-selection (Pre), UI30h, KO30h, UI60h, and WT60h (mean±SD). **(C)** Distribution of normalized read counts in all sequenced samples (median, 5-95 percentile). **(D-E)** Scatter plots displaying genes with sgRNAs found significantly enriched in cultures infected with CTL2-*cpoS*::*cat* (KO30h vs UI30h, D) or CTL2 (WT60h vs UI60h, E). Marked in green, hits with FDR ≤ 0.05 (= -log FDR ≥ 1.3) in R1 and R2; marked in blue, additional hits with FDR ≤ 0.2 (= -log FDR ≥ 0.7) in R1 and R2. Numbers mark overlapping dots.

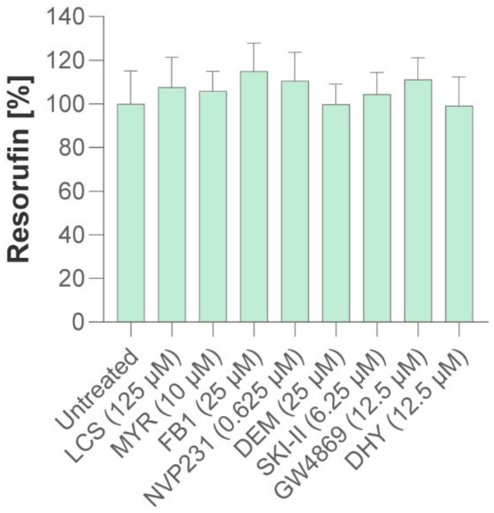


**Figure S2. The sphingolipid metabolism inhibitors were not cytotoxic at the applied concentrations.** Uninfected HeLa cells were treated with the indicated inhibitors at the indicated concentrations. Resorufin fluorescence at 25.5 hpi is displayed normalized to an untreated control (mean±SD, n=3, one-way ANOVA with Dunnett's; no significant differences compared to the untreated control).

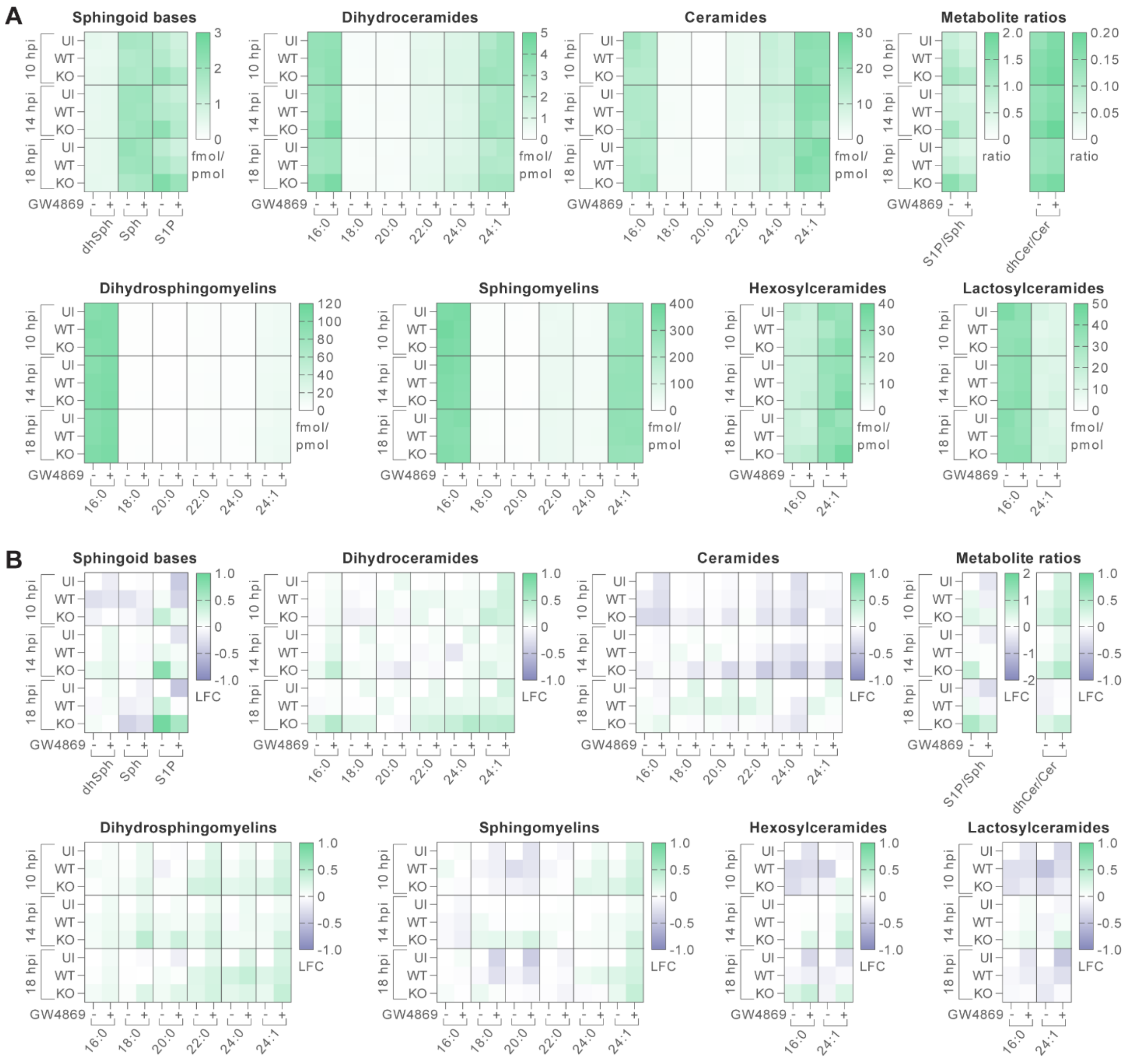


**Figure S3. CpoS deficiency caused alterations in the levels of certain sphingolipid metabolites. (A-B)** Quantification of selected sphingolipid metabolites. HeLa cells were infected with the indicated strains in the presence or absence of GW4869 (12.5 µM). At the indicated time points, cell extracts were prepared, and the indicated lipids were quantified by LC-MS/MS. Data are represented as heatmaps (means of n=3) indicating (A) metabolite levels (expressed as “fmol/pmol total sphingolipids”) or ratios or (B) log2 fold-change (LFC) of the levels or ratios compared to the respective uninfected/no inhibitor control at the same timepoint (UI, uninfected; WT, wildtype (CTL2); KO, knock-out (CTL2-*cpoS*::*cat*); Sph, sphingosine; S1P, sphingosine-1-phosphate; dhCer, dihydroceramides; Cer, ceramides).

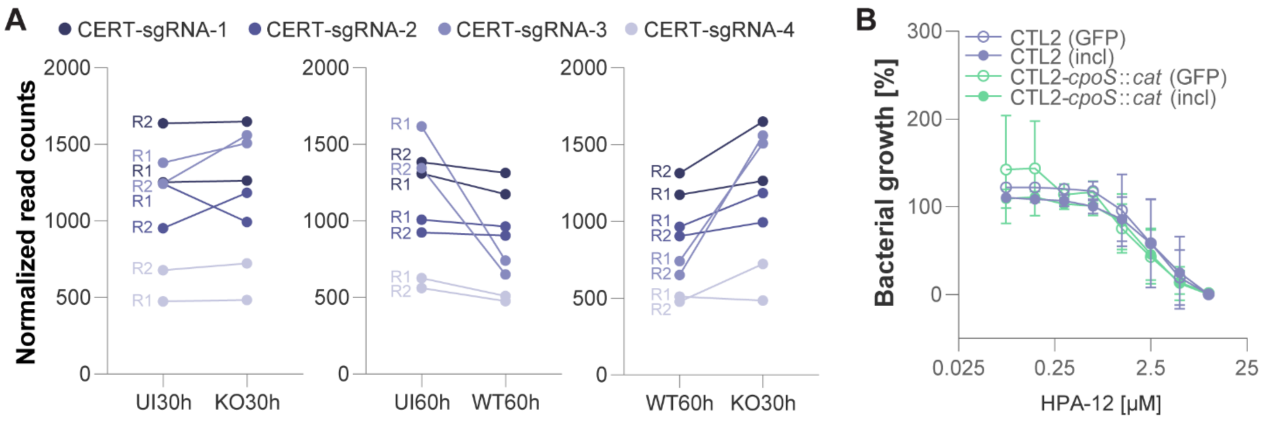


**Figure S4. Effects of CERT deficiency or inhibition on the CpoS-proficient and CpoS-deficient strains. (A)** Normalized read counts for CERT-targeting sgRNAs in the CRISPR screen. **(B)** Similar susceptibility of *cpoS* mutant and wild-type bacteria to CERT-inhibitory ceramide analog HPA-12. HeLa cells were treated with HPA-12 and parallelly infected with GFP-expressing derivatives of the indicated strains (1 IFU/cell). GFP fluorescence (GFP) and inclusion numbers (incl) at 25.5 hpi are displayed relative to values detected for the respective strains in the absence of inhibitor (mean±SD, n=3, 2-way ANOVA with Sidak's; no significant differences were observed between the two strains).

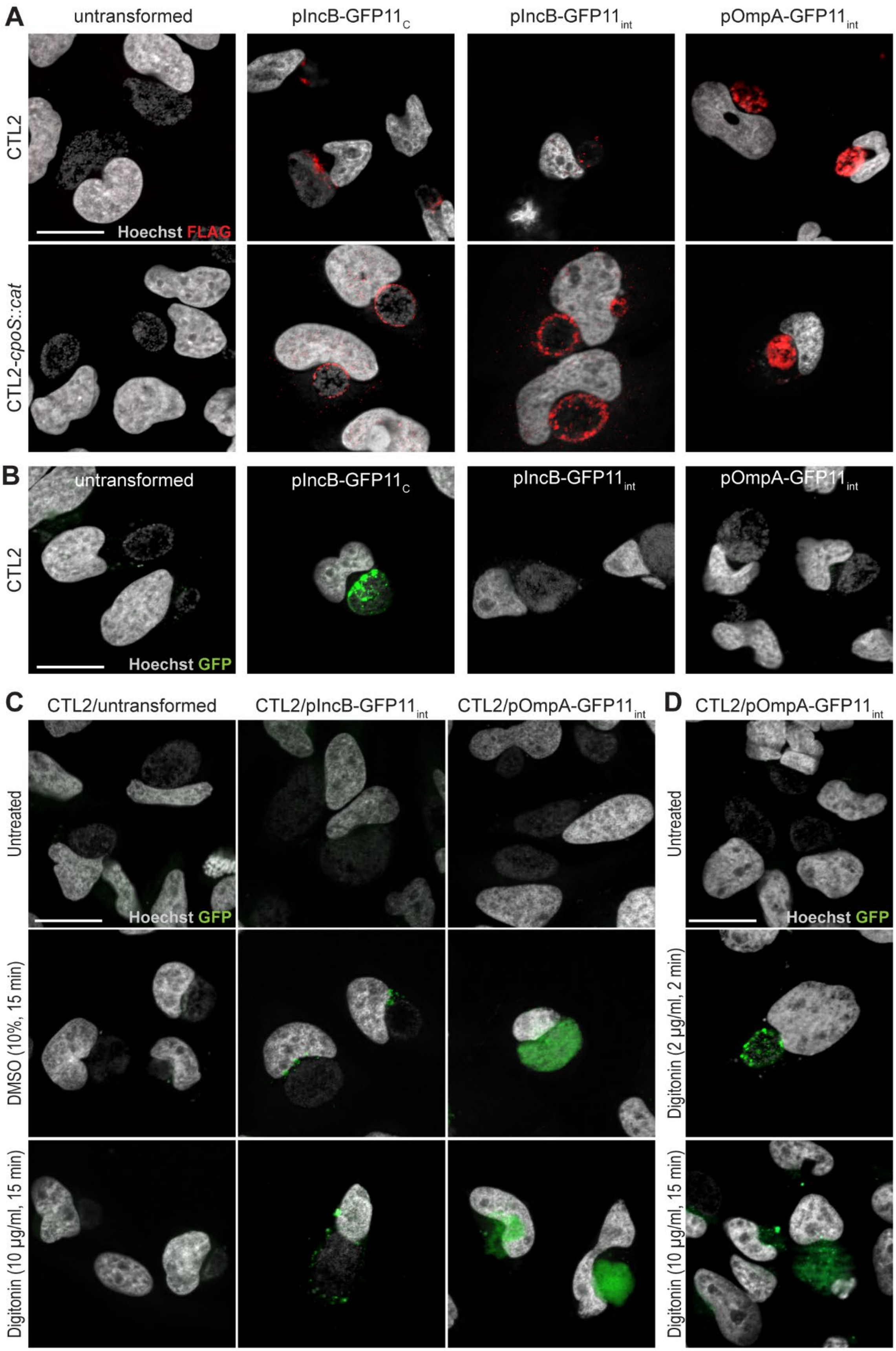
A
untransformed
pIncB-GFP11C
pIncB-GFP11int
pOmpA-GFP11int
CTL2
CTL2-cpoS::cat
Hoechst FLAG
B
untransformed
pIncB-GFP11C
pIncB-GFP11int
pOmpA-GFP11int
CTL2
Hoechst GFP
C
CTL2/untransformed
CTL2/pIncB-GFP11int
CTL2/pOmpA-GFP11int
Untreated
DMSO (10%, 15 min)
Digitonin (10 µg/ml, 15 min)
Hoechst GFP
D
CTL2/pOmpA-GFP11int
Untreated
Digitonin (2 µg/ml, 2 min)
Digitonin (10 µg/ml, 15 min)
Hoechst GFP

**Figure S5: A novel fluorescence microscopic tool enabling detection of inclusion damage. (A)** Fluorescence microscopic validation of the expression and proper localization of the GFP11-tagged constructs in CTL2 and CTL2-*cpoS*::*cat*. HeLa cells, infected with the indicated strains (5 IFU/cell), were fixed, stained, and imaged at 26 hpi (scale = 20 µm). The “donut-shaped” staining observed for OmpA-GFP11$_{int}$ aligned with its expected localization to the bacterial outer membrane. Furthermore, the patchy localization of IncB-GFP11$_{C}$ and IncB-GFP11$_{int}$ at the inclusion membrane of CTL2 was consistent with the previously reported enrichment of IncB at inclusion membrane microdomains [11,82]. In cells infected with CTL2-*cpoS*::*cat*, this patchiness was reduced, corroborating our earlier discovery that CpoS deficiency disrupts microdomain formation [11]. **(B)** Fluorescence microscopic detection of split-GFP signals in GFP1-10-expressing HeLa cells infected with the indicated strains of CTL2. HeLa cells were transfected with a plasmid driving GFP1-10 expression, infected with CTL2 (5 IFU/cell), and then fixed and imaged at 26 hpi (scale = 20 µm). **(C-D)** Detection of inclusion damage upon treatment with digitonin or DMSO. HeLa cells were transfected with a plasmid driving GFP1-10 expression and infected with the indicated strains of CTL2 (5 IFU/cell). Prior to fixation and imaging at 26 hpi, cells were treated with digitonin or DMSO (time and concentration as indicated, scale = 20 µm). A shorter duration of treatment with digitonin (as shown in (D)) preserved the morphology of bacteria in damaged inclusions.

**Methods S1. Gene blocks used for the cloning of GFP11-tagged constructs.** Gene blocks used for the generation of vector pTL2-tetO-CTL0050-GFP11x4-FLAG-CTL0050 (pOmpA-GFP11$_{int}$) and pTL2-tetO-IncB-GFP11x3-IncB-FLAG (pIncB-GFP11$_{int}$).

*Gene block (OmpA-GFP11$_{int}$)*

GCGGCCGCATGAAAAAACTCTTGAAATCGGTATTAGTGTTTGCCGCTTTGAGTTCTGCTTCCTCCTTGCAAGCTCTGC
CTGTGGGGAATCCTGCTGAACCAAGCCTTATGATCGACGGAATTCTATGGGAAGGTTTCGGCGGAGATCCTTGCGATC
CTTGCACCACTTGGTGTGACGCTATCAGCATGCGTATGGGTTACTATGGTGACTTTGTTTTCGACCGTGTTTTGCAAA
CAGATGTGAATAAAGAATTCCAAATGGGTGCCAAGCCTACAACTGCTACAGGCAATGCTGCAGCTCCATCCACTTGTA
CAGCAAGAGAGAATCCTGCTTACGGCCGACATATGCAGGATGCTGAGATGTTTACAAATGCTGCTTACATGGCATTGA
ATATTTGGGATCGTTTTGATGTATTCTGTACATTAGGAGCCACCAGTGGATATCTTAAAGGAAATTCAGCATCTTTCA
ACTTAGTTGGGTTATTCGGAGATAATGAGAACCATGCTACAGTTTCAGATAGTAAGCTTGTACCAAATATGAGCTTAG
ATCAATCTGTTGTTGAGTTGTATACAGATACTACTTTTGCTTGGAGTGCTGGAGCTCGTGCAGCTTTGTGGGAATGTG
GATGCGCGACTTTAGGCGCTTCTTTCCAATACGCTCAATCCAAGCCTAAAGTCGAAGAATTAAACGTTCTCTGTAACG
CAGCTGAGTTTACTATCAATAAGCCTAAAGGATATGTAGGGCAAGAATTCCCTCTTGATCTTAAAGCAGGAACAGATG
GTGTGACAGGAACTAAGGATGCCTCTATTGATTACCATGAATGGCAAGCAAGTTTAGCTCTCTCTTACAGACTGAATA
TGTTCACTCCCTACATTGGAGTTAAATGGTCTCGAGCAAGTTTTGATGCAGACACGATTCGTATTGCTCAGCCGAAGT
CAGCTACAACTGTCTTTGATGTTACCACTCTGAACCCAACTATTGCTGGAGGTTCGGGACGTGACCACATGGTCCTTC
ATGAGTATGTAAATGCTGCTGGGATTACAGGTGGCTCTGGAGGTAGAGATCATATGGTTCTCCACGAATACGTTAACG
CCGCAGGCATCACTGGCGGATCAGGTGGCAGGGATCACATGGTACTCCATGAATATGTGAACGCTGCTGGAATCACAG
GCGGTAGCGGCGGTCGGGACCATATGGTCTTGCACGAATATGTCAATGCTGCCGGTATCACCATGGACTACAAGGATG
ACGACGATAAGTCAGCTGGCGATGTGAAAGCTAGCGCAGAGGGTCAGCTCGGAGATACCATGCAAATCGTTTCCTTGC
AATTGAACAAGATGAAATCTAGAAAATCTTGCGGTATTGCAGTAGGAACAACTATTGTGGATGCAGACAAATACGCAG
TTACAGTTGAGACTCGCTTGATCGATGAGAGAGCTGCTCACGTAAATGCACAATTCCGCTTCTAATAAGTCGAC

*Gene block (IncB-GFP11$_{int}$)*

CGGCCGATGGTTCATTCTGTATACAATTCATTGGCTCCAGAAGGTTTTAGCCAAGTCTCTATTCAACCCAGTCAGATT
CCAACCAGCAAAAAAGTAATGATTGCGATAATGACTCTTTTTGCACTCACAGCCATTGCAGCAATAGTCCTTTCCATC
GTTACAGTTTGTGGAGGGTTTCCTTTTCTTCTTGCTGCACTTAACGGTTCGGGACGTGACCACATGGTCCTTCATGAG
TATGTAAATGCTGCTGGGATTACAGGTGGCTCTGGAGGTAGAGATCATATGGTTCTCCACGAATACGTTAACGCCGCA
GGCATCACTGGCGGATCAGGTGGCAGGGATCACATGGTACTCCATGAATATGTGAACGCTGCTGGAATCACAATGTCA
ACCGTAACTATTGGTGCATGCGTATCCTTGCCGATATTTACTTGCATAGCTACAACGTTATTACTTCTTTGTCTCCGT
AATATCGAACTCCTAGCCAGACCGCAAGTATTGACCCTCTCCACTCAATTCAGCCCAACAAAACCTCAAGAAGACTAC
AAGGATGACGACGATAAGTAATAAGTCGAC

*Legend*

Restriction sites (NotI/EagI and SalI) Start codons Genes encoding OmpA (aa 1-326 and aa 327-394) or IncB (aa 1-65 and aa 66-115) GFP1-11 repeats Linker sequences FLAG tag Stop codons
